# PEX14 functions as a molecular link between optineurin and pexophagy in human cells

**DOI:** 10.1101/2024.05.31.596776

**Authors:** Hongli Li, Suyuan Chen, Celien Lismont, Bram Vandewinkel, Mohamed A. F. Hussein, Cláudio F. Costa, Dorien Imberechts, Yiyang Liu, Jorge E Azevedo, Wim Vandenberghe, Steven Verhelst, Myriam Baes, Marc Fransen

**Affiliations:** Laboratory of Peroxisome Biology and Intracellular Communication, Department of Cellular and Molecular Medicine, KU Leuven, Leuven, Belgium; ISAS, Leibniz Institut für Analytische Wissenschaften, Dortmund, Germany; Department of Chemical Biology, Max Planck Institute of Molecular Physiology, Dortmund, Germany; Laboratory of Viral Cell Biology and Therapeutics, Department of Cellular and Molecular Medicine, KU Leuven, Leuven, Belgium; Department of Biochemistry, Faculty of Pharmacy, Assiut University, Egypt; Laboratory for Parkinson Research, Department of Neurosciences, KU Leuven, Leuven, Belgium; Laboratory for Biochemical Neuroendocrinology, Department of Human Genetics, KU Leuven, Leuven, Belgium; Instituto de Biologia Molecular e Celular (IBMC), Universidade do Porto, Porto, Portugal; Laboratory of Chemical Biology, Department of Cellular and Molecular Medicine, KU Leuven, Leuven, Belgium; Laboratory of Cell Metabolism, Department of Pharmaceutical and Pharmacological Sciences, KU Leuven, Leuven, Belgium

## Abstract

Pexophagy, the selective degradation of peroxisomes, is essential for removing excess or dysfunctional peroxisomes, and its dysregulation is linked to various diseases. Previous research has shown that optineurin (OPTN), an autophagy receptor involved in mitophagy, aggrephagy, and xenophagy, can induce pexophagy in HEK-293 cells. However, the underlying mechanism remains unclear. In this study, we used proximity labeling to identify PEX14, a peroxisomal membrane protein, as a neighboring partner of OPTN. Biochemical analyses revealed that PEX14 and OPTN interact through their respective coiled-coil and ubiquitin-binding domains. Further analyses demonstrated that the C-terminal half of overexpressed OPTN triggers pexophagy, likely by forming oligomers with endogenous OPTN. The co-localization of PEX14-OPTN complexes with LC3, combined with the suppression of OPTN-mediated peroxisome degradation by bafilomycin A1, supports a model in which PEX14 acts as a docking site for OPTN on the peroxisomal membrane, enabling the recruitment of the autophagic machinery for OPTN-mediated pexophagy.

**Summary:** This study uncovers and defines the peroxisomal membrane protein PEX14 as a key player in optineurin-driven pexophagy, advancing our mechanistic understanding of this cellular process. These findings open new avenues for developing therapeutic strategies targeting diseases associated with defective pexophagy.

## Introduction

Macroautophagy (here referred to as autophagy) is a critical cellular process wherein unnecessary or damaged intracellular components are enclosed within double-membrane vesicles, called autophagosomes, and subsequently transported to the lysosomal compartment for degradation and recycling (Klionsky et al., 2021). Acting as a vigilant guardian, autophagy plays a crucial role in maintaining cellular and organismal health by efficiently managing responses to various sources of stress (Ravanan et al., 2017). Impaired autophagy flux can result in the accumulation of dysfunctional organelles, contributing to cellular dysfunction by promoting the buildup of oxidative stress and disrupting essential signaling pathways (Rahman et al., 2023). One such organelle is the peroxisome, renowned for its pivotal role in cellular lipid and H_2_O_2_ metabolism (Wanders et al., 2023).

In mammals, the autophagic degradation of peroxisomes, known as pexophagy, can occur through different mechanisms (Li et al., 2021; Bajdzienko and Bremm, 2024). For example, a well-characterized pathway involves the recruitment of canonical autophagy receptors like NBR1 (neighbor of BRCA1 gene 1 protein) and SQSTM1/p62 (sequestosome-1) (Deosaran et al., 2013; Yamashita et al., 2014; Zhang et al., 2015; Riccio et al., 2019), which target ubiquitinated proteins (e.g., PEX5 (peroxin-5) (Zhang et al., 2015; Nordgren et al., 2015) and ABCD3 (ATP-binding cassette sub-family D member 3) (Sargent et al., 2016) at the peroxisomal membrane and bind to MAP1LC3/LC3 (microtubule-associated protein 1A/1B light chain 3B) on the autophagosomal membrane (Gubas and Dikic, 2022). In addition, under specific environmental conditions (e.g., amino acid starvation), lipidated LC3 (called LC3-II) can be recruited to the peroxisomal membrane in a non-canonical manner through direct interaction with the peroxisomal membrane protein PEX14 (peroxin-14) (Hara-Kuge and Fujiki, 2008; Jiang et al., 2015). This membrane protein, a central player in peroxisomal matrix protein import (Fransen et al., 1998), can also recruit ATG9A (autophagy-related protein 9A), a phospholipid scramblase whose activity plays a key role in autophagosomal membrane expansion (Maeda et al., 2020), through its interaction with TNKS1 (poly[ADP-ribose] polymerase tankyrase-1) and TNKS2 (poly[ADP-ribose] polymerase tankyrase-2) (Li et al., 2017). Moreover, HSPA8 (heat shock cognate 71 kDa protein) has been reported to mediate the translocation of the E3 ligase STUB1 (E3 ubiquitin-protein ligase CHIP) to oxidatively-stressed peroxisomes, leading to their turnover by autophagy (Chen et al., 2020). Furhermore, hypoxia and iron chelation have been reported to promote pexophagy by upregulating BNIP3L (BCL2/adenovirus E1B 19 kDa protein-interacting protein 3-like) through a mechanism yet to be fully characterized, bypassing ubiquitin priming (Wilhelm et al., 2022; Barone et al., 2023). Alterations in pexophagy flux can lead to diseases, either directly (Nazarko, 2017; Cho et al., 2018) or through the impairment of other selective autophagy pathways (Germain et al., 2024).

Recently, we demonstrated that ectopic expression of OPTN (optineurin), a recognized autophagy receptor involved in targeting depolarized mitochondria, protein aggregates, and intracellular bacterial pathogens (Qiu et al., 2022), induces pexophagy in HEK-293 cells (Li et al., 2023). However, the precise mechanisms underlying this process remain unclear. OPTN contains several structural domains crucial for its involvement in autophagy, including a ubiquitin-binding domain named NOA, a C-terminal ubiquitin-binding zinc finger (ZF), and an LC3-interacting region (LIR) (Korac et al., 2013; Weil et al., 2018). OPTN can also interact with diverse protein aggregates through its C-terminal coiled-coil (CC) domain, independently of ubiquitin (Korac et al., 2013). In addition, it employs an unconventional mechanism to initiate mitophagy by interacting with and activating TBK1 (TANK-binding kinase 1), a serine/threonine kinase that directly binds to the phosphoinositide 3-kinase (PI3K) complex (Nguyen et al., 2023; Yamano et al., 2024). Phosphorylation of OPTN by TBK1 increases its affinity for ubiquitin chains and LC3, thereby promoting mitophagy (Nguyen et al., 2023). Furthermore, the LIR-LC3 interaction can drive the ubiquitin-independent recruitment of OPTN, further enhancing mitophagy (Padman et al., 2019). OPTN also harbors a leucine zipper (LZ) domain capable of forming a complex with ATG9A vesicles, which initiates the formation of local autophagosomal membranes (Yamano et al., 2020). Notably, dysfunctions in OPTN-mediated autophagy have been associated with various diseases, including but not limited to amyotrophic lateral sclerosis, glaucoma, and Crohn’s disease (Qiu et al., 2022).

In this study, we identified PEX14 as a component of the OPTN interactome using proximity labeling proteomics. We also demonstrate that the interaction between OPTN and PEX14 is mediated by the NOA domain of OPTN and the CC domain of PEX14. Furthermore, we observed that PEX14-OPTN complexes co-localize with LC3 and that OPTN-mediated peroxisome degradation is inhibited by bafilomycin A1. Collectively, these findings suggest that PEX14 acts as a molecular bridge, linking OPTN to the pexophagy process.

## Results

### OPTN-GFP-mediated pexophagy displays divergent patterns across human cell lines

Using HEK-293 cells stably expressing po-mKeima, a functionally validated pH-sensitive red fluorescent reporter for pexophagy with an excitation peak that shifts from 440 nm at neutral pH to 586 nm at acidic pH and an emission peak at 620 nm, we recently discovered that (i) both endogenous and GFP-tagged OPTN can localize to peroxisomes, and (ii) the expression of OPTN-GFP is sufficient to induce pexophagy, even under basal growth conditions (Li et al., 2023). At the outset of this study, we observed that expression of OPTN-GFP in other cell lines stably expressing po-mKeima induced pexophagy to varying extents (Fig. S1). This variability was correlated with the recruitment of OPTN-GFP to peroxisomes, as evidenced by its colocalization with the peroxisomal marker mCherry-PTS1 and a marked reduction in peroxisome numbers in HEK-293 and HCT116 cells, but not in HeLa cells (Fig. 1A, left panels; see panel 1D for comparison). In addition, co-expression of OPTN-GFP with a high-affinity, plasma membrane-targeted anti-GFP nanobody resulted in the redistribution of peroxisomes to the plasma membrane in HEK-293 and HCT116 cells, while no such effect was observed in HeLa cells (Fig. 1A, right panels). Notably, the expression of OPTN-GFP in HEK-293 or HCT116 cells consistently led to the formation of clusters containing OPTN-GFP and peroxisomes (Fig. 1B), which were more clearly visualized using structured illumination microscopy (Fig. 1C). These clusters could be labeled with the autophagy marker GFP-LC3 (Fig. 2). Furthermore, no co-clustering of OPTN-GFP with mitochondria (Fig. S2A), the endoplasmic reticulum (Fig. S2A), or lysosomes (Fig. S2B) was observed, indicating that the co-clustering of OPTN-GFP with organelles is peroxisome-specific. Occasionally, similar clusters containing endogenously expressed OPTN and peroxisomes were also observed (Fig. 1D). Importantly, while OPTN-mediated degradation of peroxisomes was notably inhibited by the pharmacological autophagy inhibitor bafilomycin A1 (Fig. S3A,B), overexpression of OPTN-GFP did not induce general autophagy, as evidenced by Western blot analyses showing unchanged LC3B and SQSTM1 protein levels (Fig. S3C,D). Collectively, these findings demonstrate the recruitment of OPTN to peroxisomes and highlight its role in promoting pexophagy across different human cell types.

**Figure 1.**
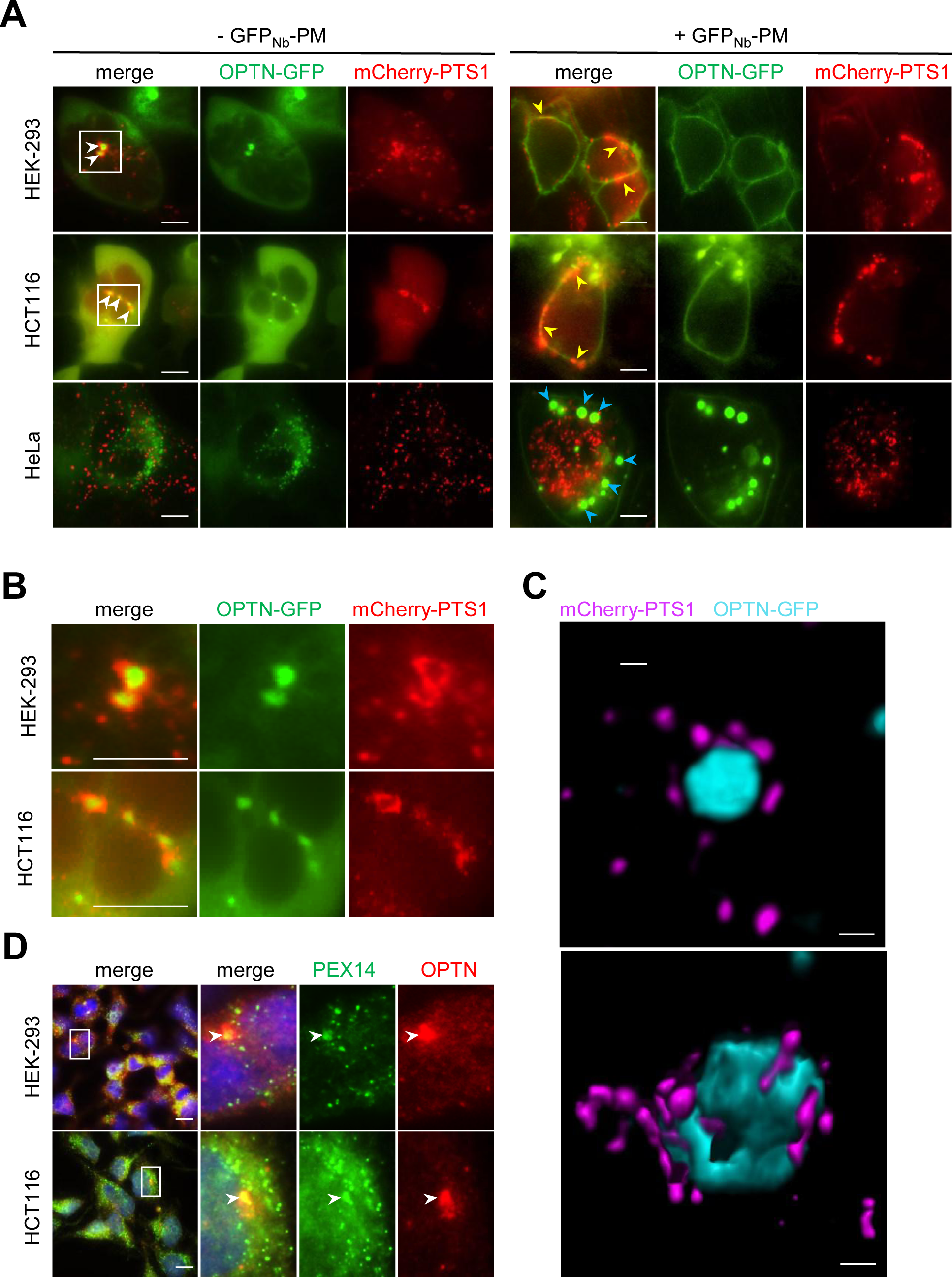
Cell type-specific recruitment of optineurin to peroxisomes. **(A)** Distribution patterns of OPTN-GFP and mCherry-PTS1 in three distinct cell lines, either co-expressing (right panels) or not (left panels) a high-affinity plasma membrane-targeted anti-GFP nanobody (GFP_Nb_-PM). Before analysis, the cells were incubated for 2 h with 200 nM BafA1. Representative images are shown. The white arrowheads mark the colocalization of OPTN-GFP and mCherry-PTS1. The yellow arrowheads highlight positions where peroxisomes are associated with the plasma membrane. The blue arrowheads indicate GFP_Nb_-PM-induced foci which may represent OPTN-GFP condensates that assemble in a concentration-dependent manner (López-Palacios and Andersen, 2023; Yang et al., 2024). **(B)** Enlargements of the boxed areas in panel a. **(C)** High-resolution imaging and volume rendering of two mCherry-PTS1/OPTN-GFP aggregates in HEK-293 cells. **(D)** Distribution patterns of endogenous PEX14 and OPTN in HEK-293 and HCT116 cells. Scale bars: **A, B,** and **D**, 10 μm; **C**, 500 nm.

**Figure 2.**
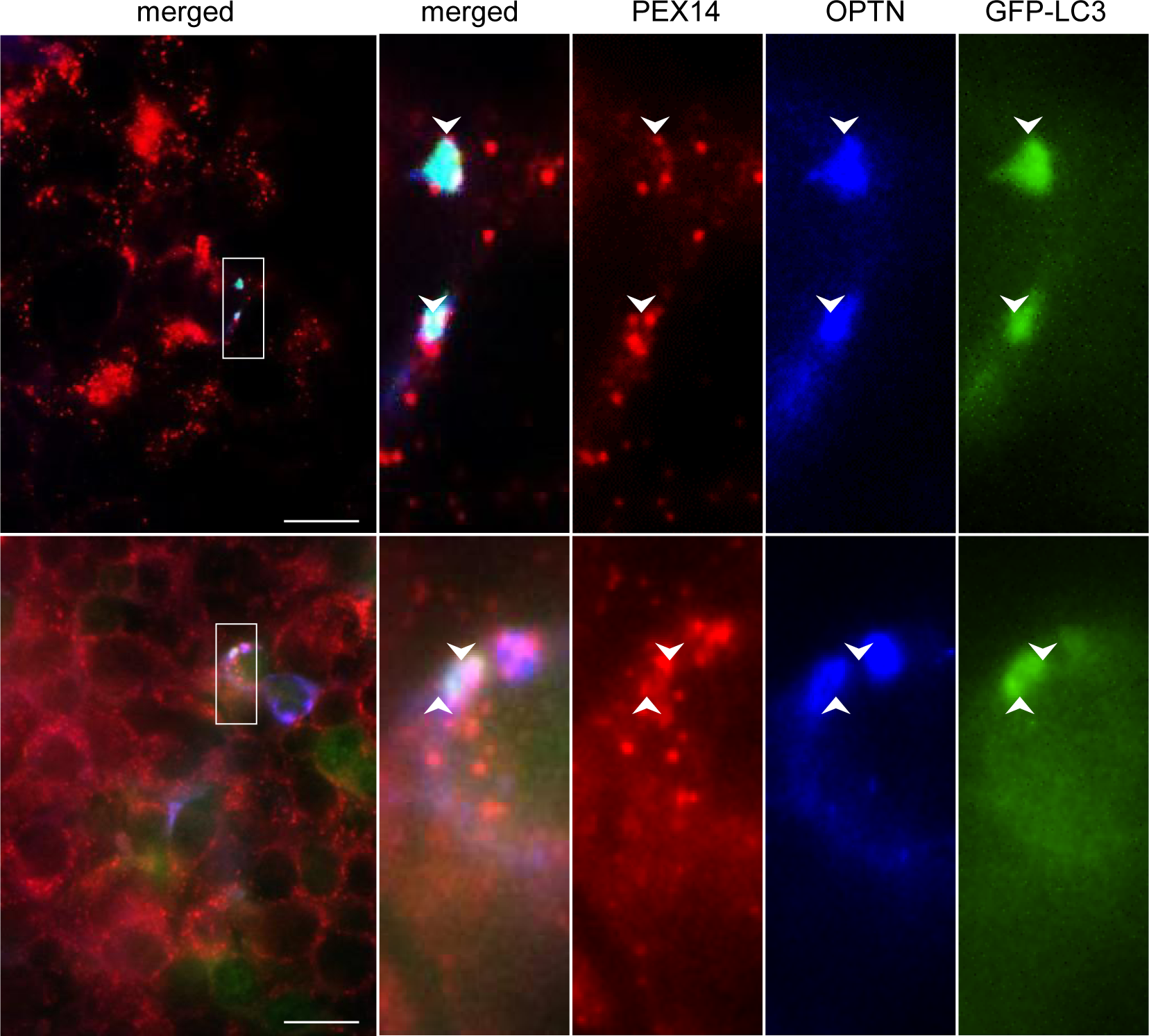
GFP-LC3 localizes to PEX14/OPTN-positive structures in HEK-293 cells. The cells were transfected with GFP-LC3-encoding plasmid and, after 2 days, they were fixed and processed for immunofluorescence microscopy. A mouse antiserum against PEX14 and a rabbit antiserum against OPTN were used for labeling. White arrowheads indicate sites where GFP-LC3 colocalizes with OPTN and the peroxisomal marker PEX14. Enlarged views of the boxed areas are shown on the right. Scale bar, 10 µm.

### The peroxisomal membrane protein PEX14 is a proximity partner of OPTN

To gain a better understanding of the molecular factors involved in OPTN-mediated pexophagy, we conducted a series of proximity-dependent biotin identification (BioID) experiments in HEK-293 cells transiently expressing OPTN-GFP fused to the miniTurbo biotin ligase (Fig. 3A). Upon the addition of biotin to the medium, this fusion protein enables the proximity-dependent covalent biotin labeling of proteins within a 10 nm radius (Qin et al., 2021). Subsequently, through streptavidin-based affinity purification of the biotinylated proteins combined with mass spectrometry identification, the neighboring proteins of OPTN can be identified at the molecular level. Among the top-ranked proteins identified in three independent replicates were TBK1, a well-known interaction partner of OPTN (Morton et al., 2008), and the peroxisomal membrane protein PEX14 (Fig. 3B). Notably, no other peroxisomal protein exhibited the same enrichment factor as PEX14, which led us to hypothesize that PEX14 and OPTN may form a protein complex in these cells.

**Figure 3.**
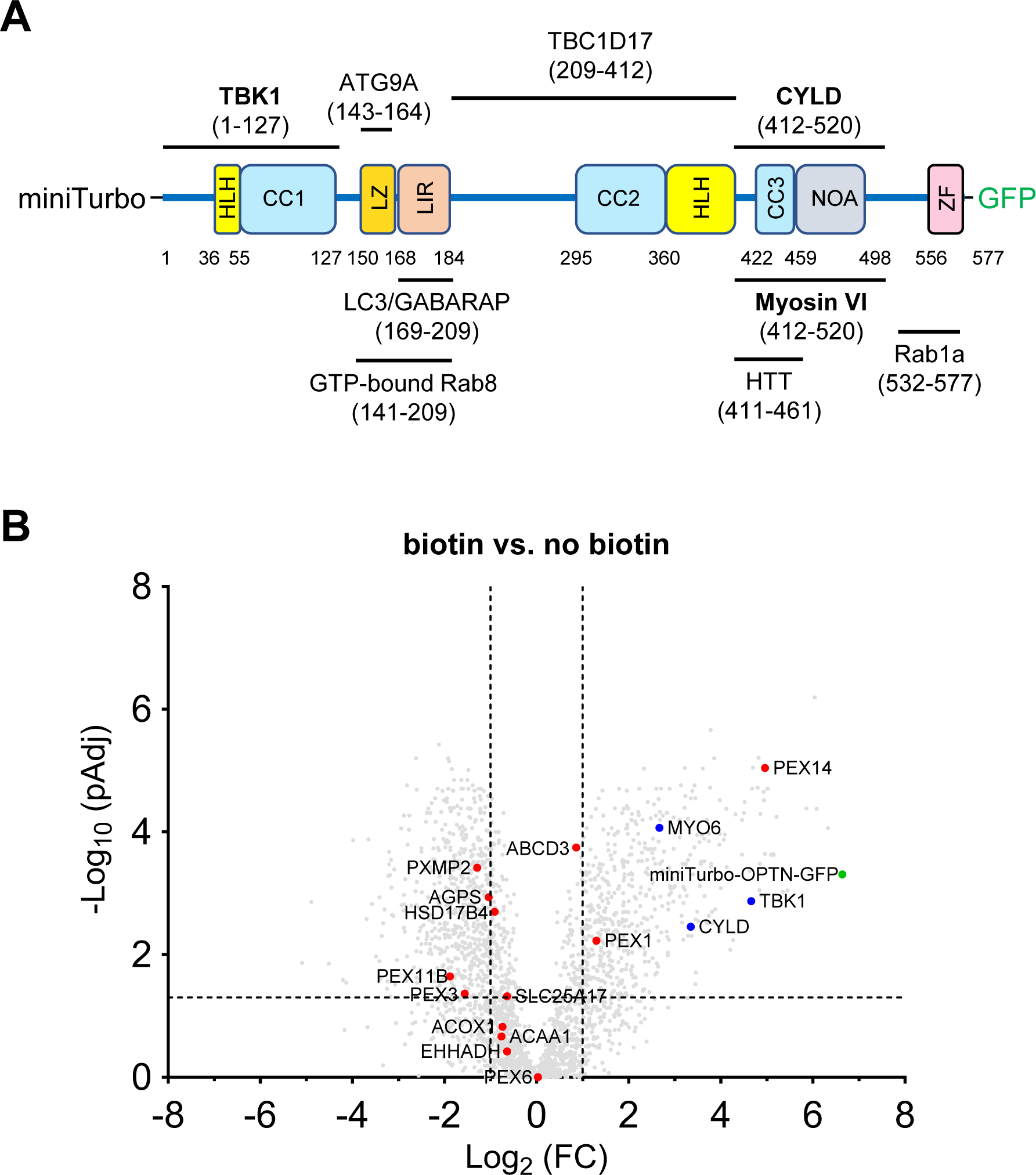
PEX14 is a proximity partner of OPTN. **(A)** Schematic representation of the distinct domains and major interactions of the miniTurbo-OPTN-GFP fusion protein. ATG9A, autophagy-related protein 9A; CC, coiled-coil; CYLD, cylindromatosis; HLH, helix-loop-helix; HTT, huntingtin; LZ, leucine zipper domain; LC3/GABARAP, microtubule-associated protein 1A/1B-light chain 3/γ-aminobutyric acid receptor-associated protein; LIR, LC3-interacting region; Rab, Ras-related protein; TBC1D17, TBC1 domain family member 17; TBK1, TANK (TRAF-associated NF-kB activator) binding kinase 1; NOA, NEMO-OPTN-ABIN domain; ZF, zinc finger. **(B)** Volcano plot displaying the Log_2_-fold changes (FC) in protein abundance alongside the corresponding Log_10_-fold adjusted p-values (pAdj) for 2856 proteins obtained through streptavidin-affinity purification and identification/quantification by LC–MS/MS. The analysis was performed on miniTurbo-OPTN-GFP-expressing HEK-293 cells (n = 4 biological replicates), assayed under conditions with or without 50 µM biotin. Green, blue, and red dots represent miniTurbo-OPTN-GFP, known OPTN interaction partners, and bona fide peroxisomal proteins that could be identified in the affinity-purified fractions. Vertical and horizontal dotted lines denote the FC (2x) and pAdj (0.05) cut-offs, respectively.

To test this possibility, we performed GFP-Trap analyses in HEK-293 cells overexpressing OPTN using three methods: transient expression by electroporating cells with an OPTN-encoding mammalian expression plasmid, inducible expression in OPTN-GFP Flp-In T-REx 293 cells treated with doxycycline, and constitutive expression by infecting Flp-In T-REx 293 cells with lentivirus. In some experiments, OPTN-GFP was expressed alongside ectopically expressed PEX14. As shown in Fig. S4A,B, both ectopically expressed and endogenous PEX14 co-immunoprecipitated with OPTN-GFP, indicating that the two proteins form a complex in these cells. Note that the expression levels of OPTN-GFP were approximately 15 times higher than those of endogenous OPTN (Fig. S4C,D).

### The C-terminal half of OPTN is sufficient to bind PEX14 and initiate pexophagy

Optineurin consists of multiple domains that either interact with various types of cargo or regulate the engulfment of cargo into autophagosomes (Fig. 3A). To gain insights into the OPTN domains responsible for interacting with PEX14 and inducing pexophagy, we co-expressed the N-terminal or C-terminal halves of OPTN, fused to GFP, with PEX14 in HEK-293 cells and conducted GFP-Trap analyses and flow cytometry-based pexophagy assays.

From these experiments, it is evident that the C-terminal 361 amino acids of OPTN, which include the NOA and ZF ubiquitin-binding domains, are sufficient for binding to PEX14 (Fig. 4A,B) and facilitating pexophagy (Fig. 4D,E). In contrast, the N-terminal 209 amino acids of OPTN, containing the TBK1-, LC3-, and ATG9A-binding domains, show weak binding to PEX14 (Fig. 4A,B) and are insufficient to induce pexophagy (Fig. 4D,E).

**Figure 4.**
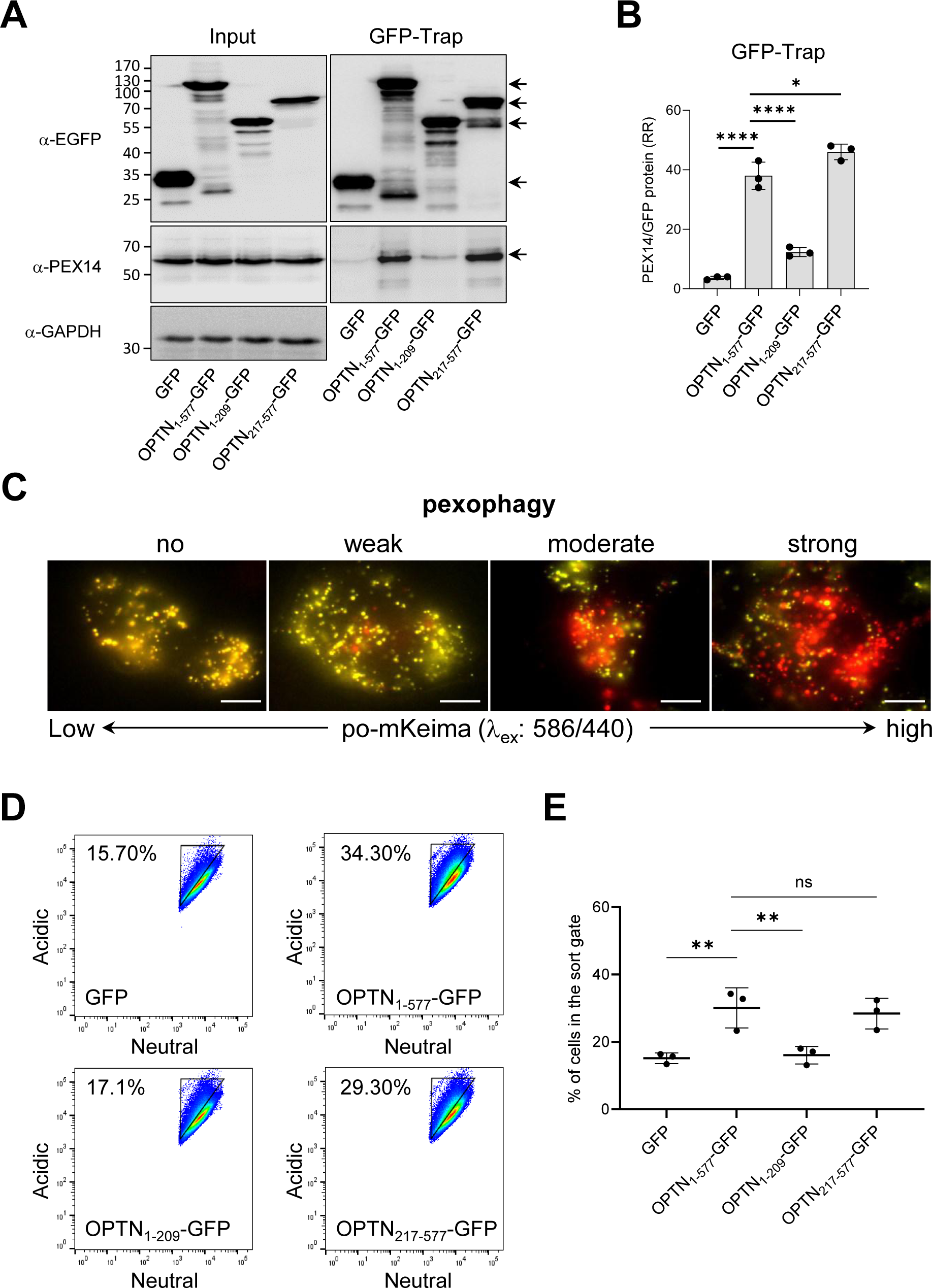
Mapping of PEX14-interacting and pexophagy-inducing regions of OPTN. Flp-In T-REx 293 cells stably expressing po-mKeima were transfected with a plasmid encoding the indicated GFP-fusion protein and cultured in regular DMEM medium. After two days, the cells were processed for GFP-Trap and FACS analyses. Pexophagy was measured by gating single-cell GFP-positive populations for decreases in fluorescence intensity in the neutral channel. **(A)** Samples of protein extracts (input) and the GFP Trap were processed for SDS-PAGE followed by immunoblotting using antibodies to GFP, PEX14, and GAPDH. Representative blots are shown. Specific protein bands are marked by arrows. Molecular mass markers (in kDa) are indicated on the left. **(B)** Densitometry quantifications of the relative ratios (RR) of PEX14/GFP and PEX14/OPTN-GFP fusion proteins retained on the GFP-Trap affinity matrix. The total signal intensities of the GFP proteins and PEX14 were both normalized to 100%. Bars represent the mean ± the standard deviation of 3 biological replicates. **(C)** Examples of po-mKeima-expressing cells displaying no, weak, moderate, or excessively high levels of pexophagy. The visuals depict overlay images of po-mKeima excited at 440 nm (false color: green) or 586 nm (false color: red). Yellowish (=low 586/440 excitation peak ratio) and reddish (=high 586/440 excitation peak ratio) dots represent peroxisomes and pexolysosomes, respectively [25]. Scale bar, 10 µm. **(D)** Representative flow cytometry plots of each group (n = 3). The different colors represent the cell density at a given position. **(E)** Quantification of the percentage of cells in the gated area. The data are shown as the mean ± SD and represent the values of 3 independent biological replicates. All conditions were statistically compared to the OPTN_1-577_-GFP condition (ns, non-significant; *, p < 0.05; **, p < 0.01; ****, p < 0.0001).

### OPTN binds to PEX14 and initiates pexophagy through its NOA domain

To delineate the precise C-terminal domain of OPTN responsible for its interaction with PEX14, we first co-expressed OPTN-GFP fusion proteins with internal deletion of CC2 (OPTN_Δ295-360_-GFP), CC3 (OPTN_Δ422-454_-GFP), NOA (OPTN_Δ454-520_-GFP), or the ZF (OPTN_Δ545-577_-GFP) domain, alongside non-tagged full-length PEX14. Results indicate that the deletion of amino acids 454 to 520, encompassing the full NOA domain, significantly diminishes the affinity of OPTN for PEX14 (Fig. 5A,B). Unfortunately, due to the very low expression levels of OPTN_545-577_-GFP (data not shown), no reliable conclusions could be drawn regarding the potential role of the ZF domain of OPTN in PEX14 binding.

**Figure 5.**
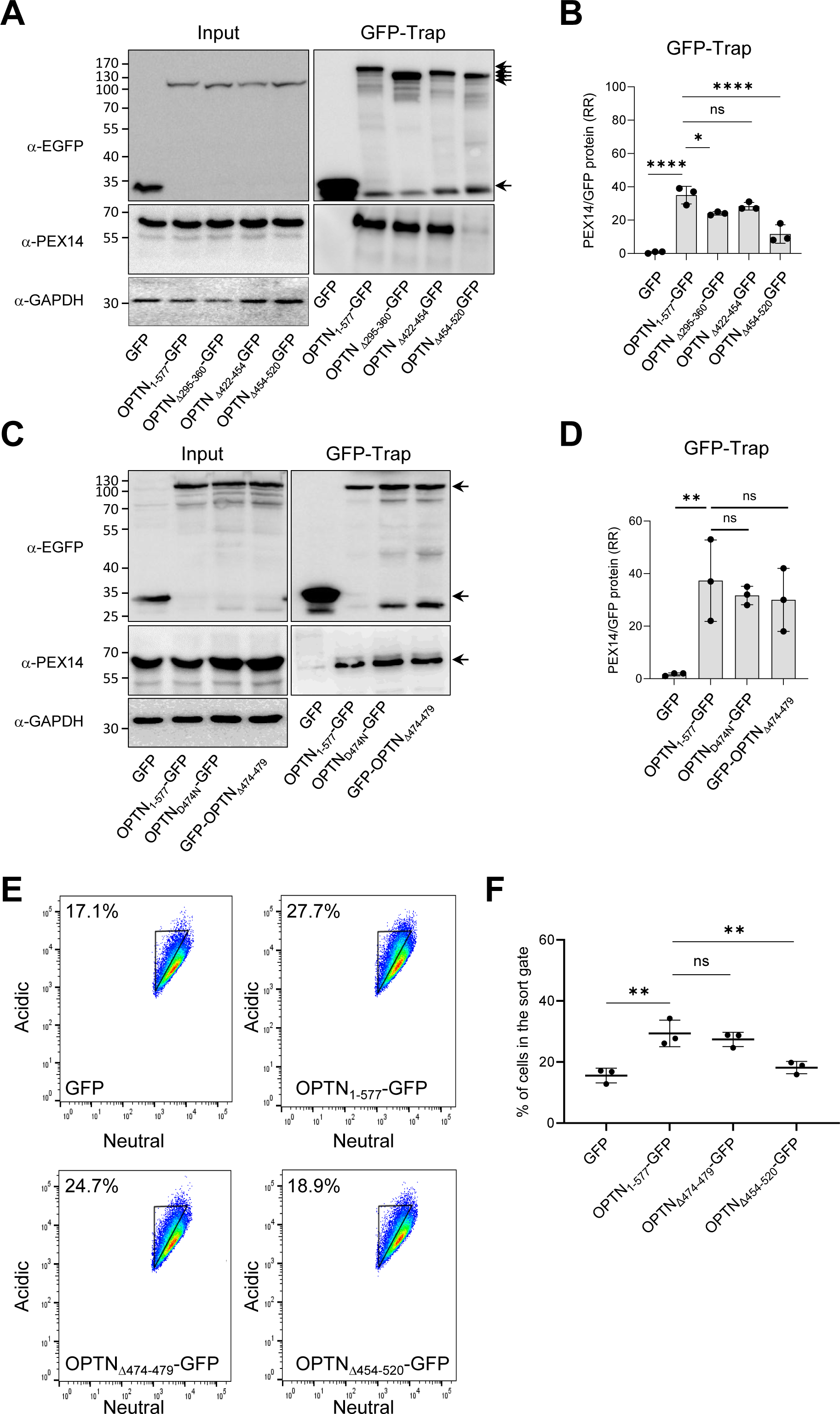
OPTN-GFP interacts with PEX14 and triggers pexophagy in a NOA-dependent manner. Flp-In T-REx 293 cells stably expressing po-mKeima were transfected with a plasmid encoding the indicated GFP-fusion protein and cultured in regular DMEM medium. After two days, the cells were processed for GFP-Trap and FACS analyses. Pexophagy was measured by gating single-cell GFP-positive populations for decreases in fluorescence intensity in the neutral channel. **(A,C)** Samples of protein extracts (input) and the GFP-Trap were processed for SDS-PAGE followed by immunoblotting using antibodies to GFP, PEX14, and GAPDH. Representative blots are shown. Specific protein bands are marked by arrows. Molecular mass markers (in kDa) are indicated on the left. **(B,D)** Densitometry quantifications of the RR of PEX14/GFP and PEX14/OPTN-GFP fusion proteins retained on the GFP-Trap affinity matrix. The total signal intensities of the GFP proteins and PEX14 were both normalized to 100%. Bars represent the mean ± the standard deviation of 3 biological replicates. **(E)** Representative flow cytometry plots of each group (n = 3). The different colors represent the cell density at a given position. **(F)** Quantification of the percentage of cells in the gated area. The data are shown as the mean ± SD and represent the values of 3 independent biological replicates. All conditions were statistically compared to the OPTN_1-577_-GFP condition (ns, non-significant; *, p < 0.05; **, p < 0.01; ****, p < 0.0001).

We then investigated whether the ubiquitin-binding capacity of OPTN’s NOA domain is necessary for its interaction with PEX14. We co-expressed PEX14 with OPTN_D474N_-GFP or GFP-OPTN_Δ474-479_, two OPTN variants in which key residues in the NOA ubiquitin-binding domain were either mutated or deleted (Gleason et al., 2011; Wild et al., 2011). These experiments show that ubiquitin binding in the NOA domain of OPTN is not required for the OPTN-PEX14 interaction (Fig. 5C,D). In addition, OPTN variants that can bind PEX14 are also capable of inducing pexophagy (Fig. 5E,F).

### Knockdown of endogenous OPTN disrupts OPTN_217-577_-GFP-mediated pexophagy

It is known that the LC3 and ATG9A-binding domains of OPTN are important for facilitating autophagosome formation and participating in selective autophagy (Zhao et al., 2023).

However, as shown in Fig. 4D,E, a truncated OPTN variant lacking these domains (OPTN_217-577_-GFP) is still capable of inducing pexophagy. Given that OPTN primarily exists as a trimer in vivo (Gao et al., 2014), we wondered whether the truncated OPTN-GFP protein might oligomerize with endogenous OPTN. Therefore, we downregulated the expression levels of endogenous OPTN by transfecting HEK-293 cells with a DsiRNA targeting exon 5, encoding amino acids 57 to 123 (Fig. S5). Although we were unable to confirm by immunoprecipitation that OPTN_217-577_-GFP interacts with the endogenous protein, due to both proteins exhibiting the same migration pattern on SDS-PAGE (Fig. S5C), these experiments demonstrated that knocking down endogenous OPTN expression interferes with OPTN_217-577_-GFP-mediated pexophagy (Fig. 6).

**Figure 6.**
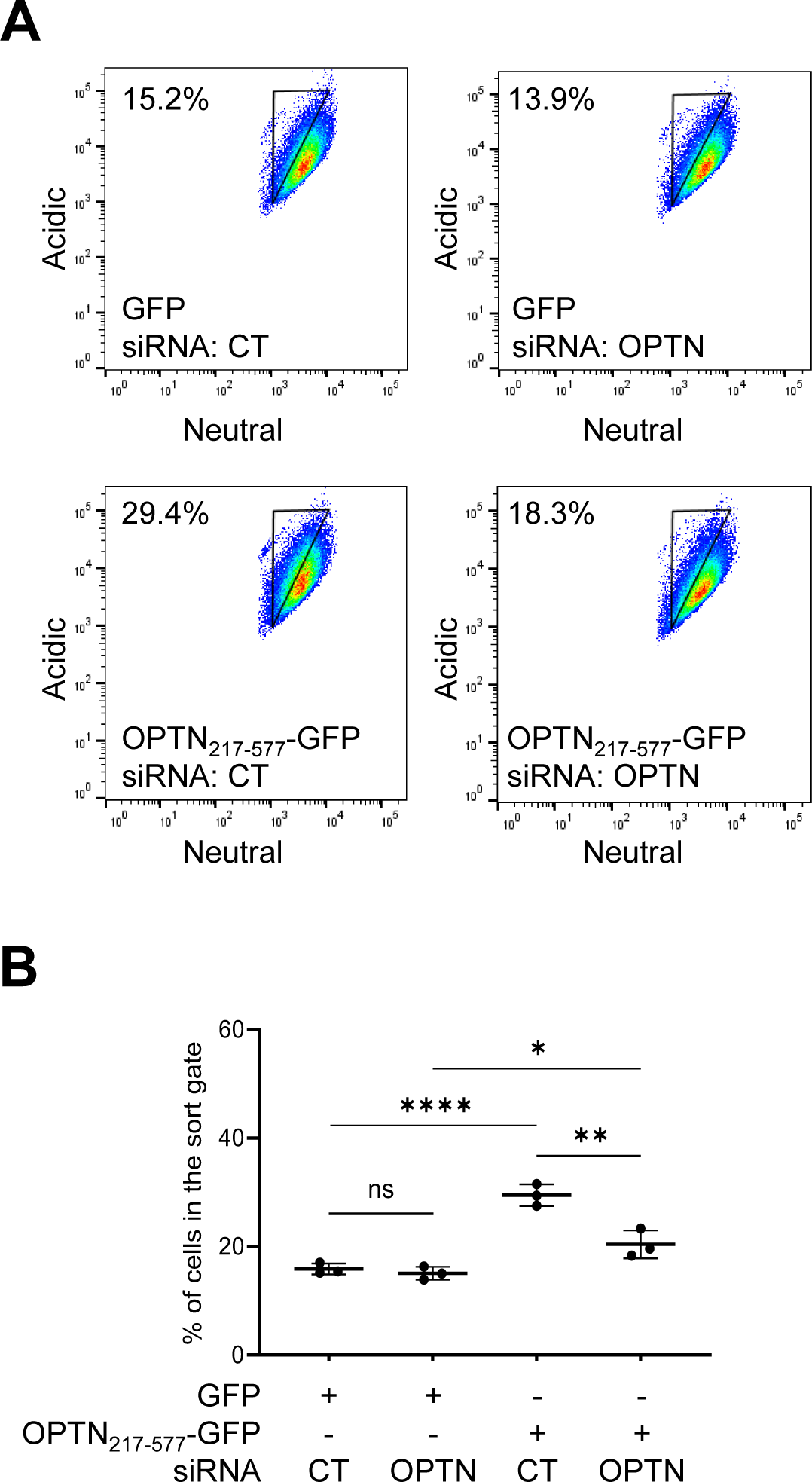
OPTN_217-577_-GFP-induced pexophagy in HEK-293 cells relies on endogenous OPTN. Flp-In T-REx 293 cells, stably expressing po-mKeima, were transfected with either a control (CT) or OPTN DsiRNA and seeded in 6-well plates. On day 2, the cells were transfected with either a GFP- or OPTN_217-577_-GFP-encoding plasmid. On day 3, the cells were transfected again with either DsiRNA CT or DsiRNA OPTN. On day 4, the cells were harvested and processed for FACS analysis. Pexophagy was measured by gating single-cell GFP-positive populations for decreases in fluorescence intensity in the neutral channel. **(A)** Representative flow cytometry plots of each group (n = 3). The different colors represent the cell density at a given position. **(B)** Quantification of the percentage of cells in the gated area. The data are shown as the mean ± SD and represent the values of 3 independent biological replicates. The impact of OPTN downregulation and OPTN_217-577_-GFP overexpression were statistically compared (ns, non-significant; *, p < 0.05; **, p < 0.01;****, p < 0.0001).

### PEX14 interacts with OPTN via its predicted coiled-coil domain

To delineate the domain on PEX14 responsible for its interaction with OPTN, we conducted GFP-Trap assays on lysates derived from HEK-293 cells co-expressing non-tagged OPTN and various PEX14-GFP variants that impair the protein’s N-terminal PEX5/PEX13/PEX19-binding domain (PEX14_65-377_-GFP), membrane anchoring region (PEX14_Δ110-126_-GFP), PEX14-binding coiled-coil domain (PEX14_Δ139-199_-GFP), or C-terminal D/E-rich region (PEX14_1-313_-GFP) (Fransen et al., 2002). These experiments demonstrate that the coiled-coil domain of PEX14 is essential for its binding to OPTN (Fig. 7A,B). In line with this, co-expression of non-tagged full-length OPTN with PEX14_1-377_-GFP, but not PEX14_Δ139-199_-GFP, promoted OPTN-induced pexophagy not only in HEK-293 cells (Fig. 7C-F) but, remarkably, also in HeLa cells (Fig. 8; see also discussion).

**Figure 7.**
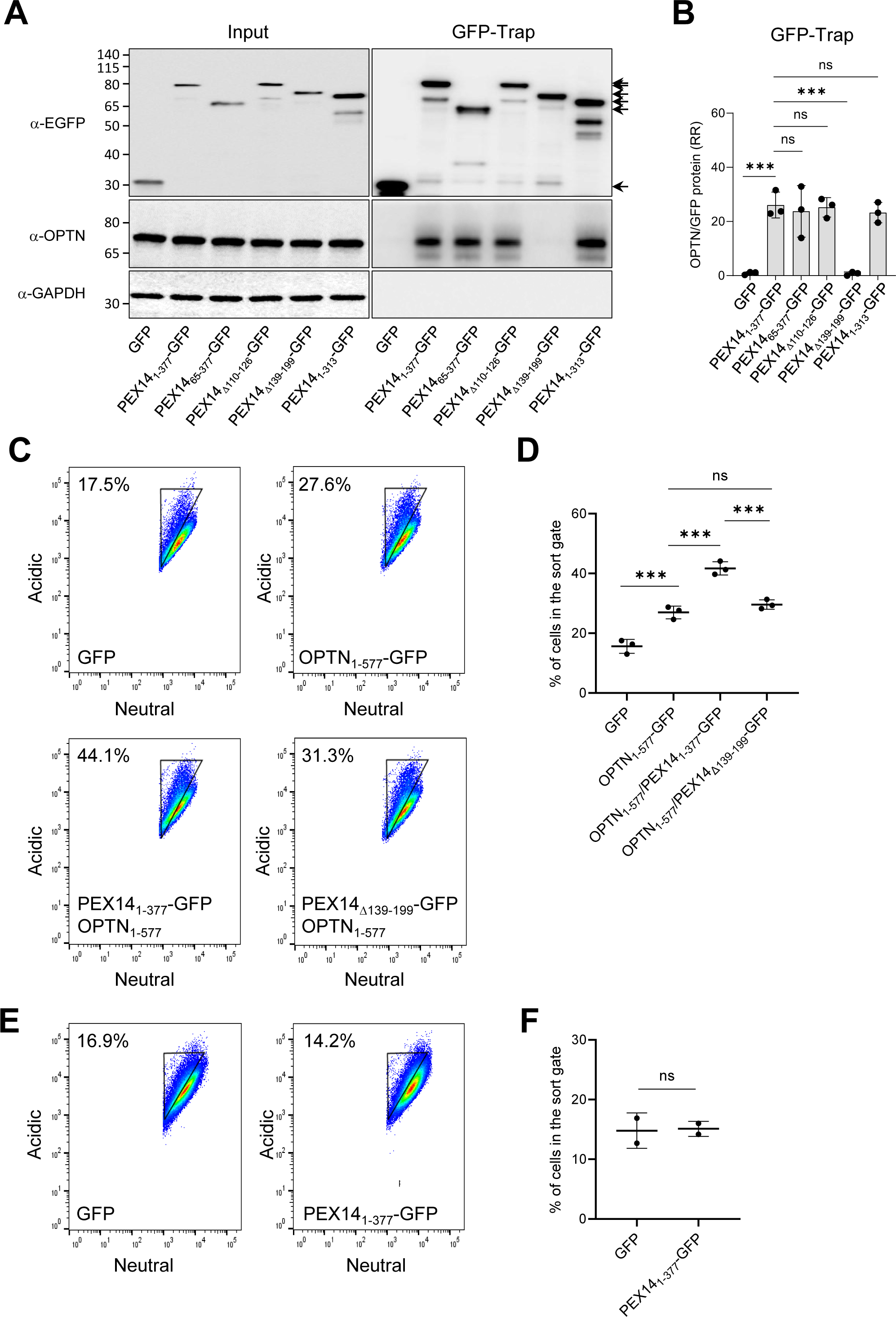
Mapping of the OPTN-interacting region of PEX14 and assessment of its expression levels on OPTN-mediated pexophagy. Flp-In T-REx 293 cells stably expressing po-mKeima were (co-)transfected with (a) plasmid(s) encoding the indicated protein(s) and cultured in regular DMEM medium. After two days, the cells were processed for GFP-Trap and FACS analysis. Pexophagy was measured by gating single-cell GFP-positive populations for decreases in fluorescence intensity in the neutral channel. **(A)** Samples of protein extracts (input) and the GFP-Trap were processed for SDS-PAGE followed by immunoblotting using antibodies to GFP, OPTN, and GAPDH. Representative blots are shown. Specific protein bands are marked by arrows. Molecular mass markers (in kDa) are indicated on the left. **(B)** Densitometry quantifications of the RR of OPTN/GFP and OPTN/PEX14-GFP fusion proteins retained on the GFP-Trap affinity matrix. The total signal intensities of the GFP proteins and PEX14 were both normalized to 100%. Bars represent the mean ± the standard deviation of 3 biological replicates. **(C,E)** Representative flow cytometry plots of each group (n = 2-3). The different colors represent the cell density at a given position. **(D,F)** Quantification of the percentage of cells in the gated area. The data are shown as the mean ± SD and represent the values of 2-3 independent biological replicates. Relevant conditions were statistically compared (ns, non-significant; ***, p < 0.001).

**Figure 8.**
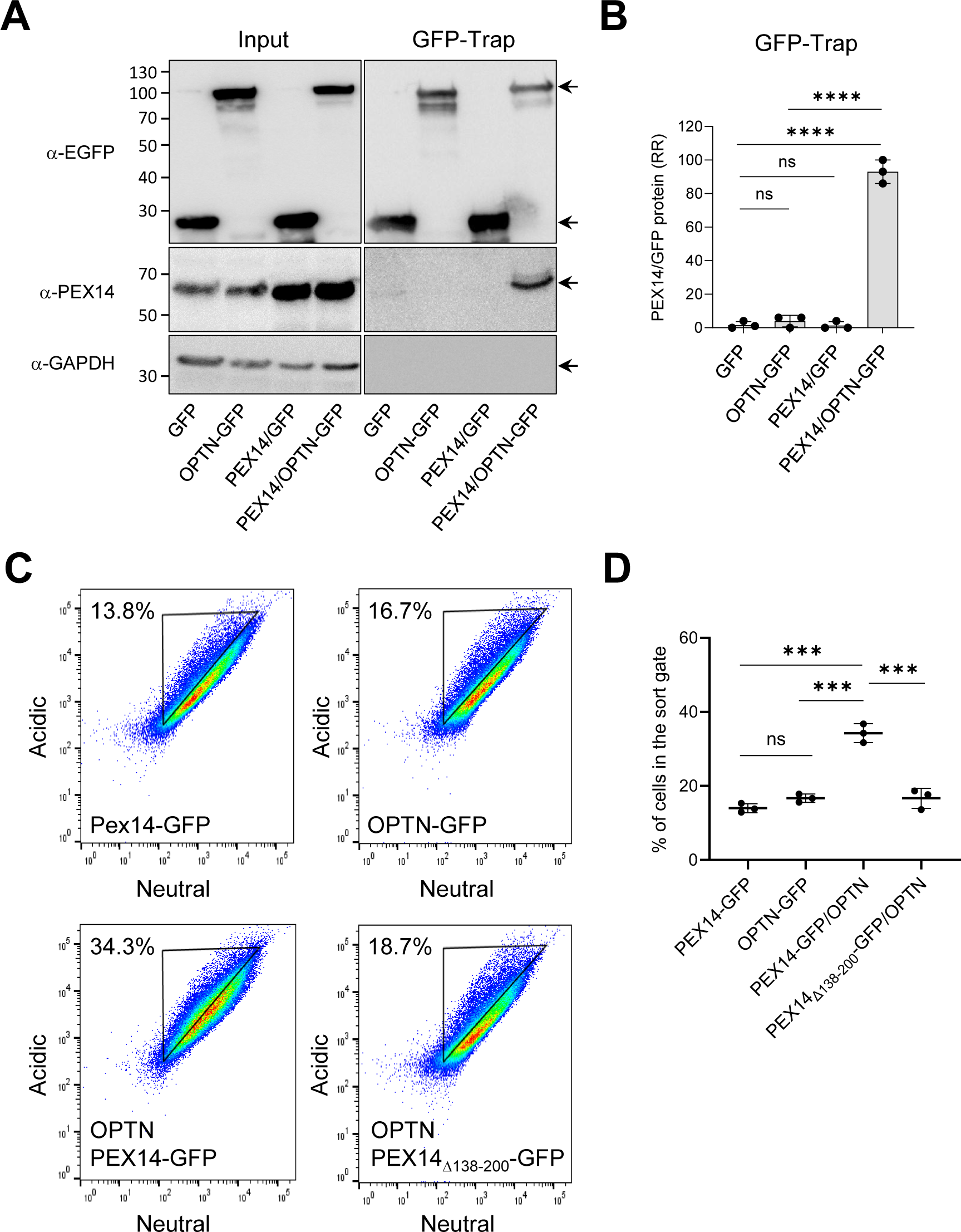
Overexpression of PEX14 induces its interaction with OPTN-GFP and triggers pexophagy in HeLa cells. HeLa cells stably expressing po-mKeima were transfected with plasmids encoding the indicated proteins and cultured in DMEM. After two days, the cells were processed for GFP-Trap and FACS analyses. Pexophagy was measured by gating single-cell GFP-positive populations for decreases in fluorescence intensity in the neutral channel. **(A)** Protein extracts (input) and the GFP-Trap were subjected to SDS-PAGE and immunoblotting with antibodies against GFP, PEX14, and GAPDH. Representative blots are shown. Specific protein bands are marked by arrows. Molecular mass markers (in kDa) are indicated on the left. **(B)** Densitometry quantifications of the RR of PEX14/GFP and PEX14/OPTN-GFP fusion proteins retained on the GFP-Trap affinity matrix. Signal intensities for GFP proteins and PEX14 were normalized to 100%. Bars represent the mean ± the standard deviation of 3 biological replicates. **(C)** Representative flow cytometry plots of each group (n = 3). The different colors represent the cell density at a given position. **(D)** Quantification of the percentage of cells in the gated area. The data are shown as the mean ± SD and represent the values of 3 independent biological replicates. Relevant conditions were statistically compared (ns, non-significant; ***, p < 0.001; ****, p < 0.0001).

### Torin-1 induces pexophagy in the absence of classical autophagy receptors

The finding that OPTN-GFP does not trigger pexophagy in HeLa cells unless PEX14 is also overexpressed (Fig. 8) led us to hypothesize that pexophagy in these cells may be mediated by one or more other classical autophagy receptors involved in pexophagy (Deosaran et al., 2013; Yamashita et al., 2014; Zhang et al., 2015; Riccio et al., 2019). To identify those receptor(s), we first examined peroxisome abundance in Hela cells lacking the five classical autophagy receptors: SQSTM1, NBR1, OPTN, NDP52, and TAX1BP1 (Lazarou et al., 2015). Surprisingly, immunofluorescence and immunoblotting analyses revealed that peroxisome abundance remained unchanged in these cells (Fig. 9). Remarkably, we found that Torin-1, a potent autophagy activator, could still induce pexophagy in po-mKeima Penta-KO HeLa cells (Fig. 10), supporting the existence of non-canonical pexophagy pathways (Hara-Kuge and Fujiki, 2008; Jiang et al., 2015; Li et al., 2017; Wilhelm et al., 2022; Barone et al., 2023).

**Figure 9.**
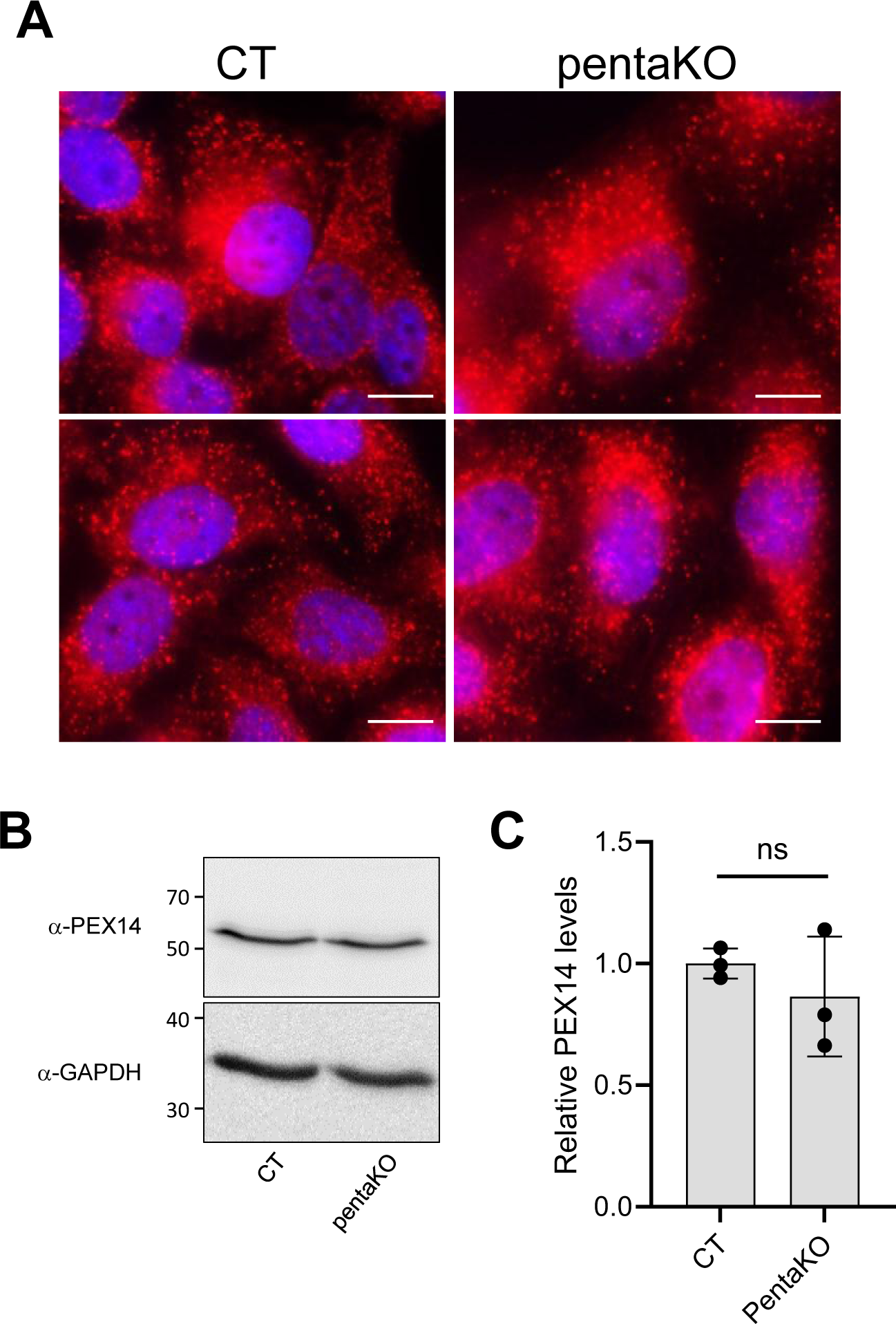
Peroxisome abundance remains unchanged in Hela cells lacking classical autophagy receptors. Control (CT) and PentaKO HeLa cells were analyzed by immunofluorescence and immunoblotting using PEX14-specific antibodies. **(A)** Representative images showing peroxisomes in both CT and pentaKO cells, with nuclei counterstained by DAPI. Scale bar, 10 µm. **(B)** Representative immunoblot illustrating the relative abundance of PEX14 in CT and pentaKO cells. **(C)** Densitometry quantification of PEX14 levels in CT and pentaKO cells, with PEX14 signal intensities normalized to the mean value of the CT cells. Data represent the mean ± the standard deviation of 3 biological replicates. No significant differences (ns) were observed.

**Figure 10.**
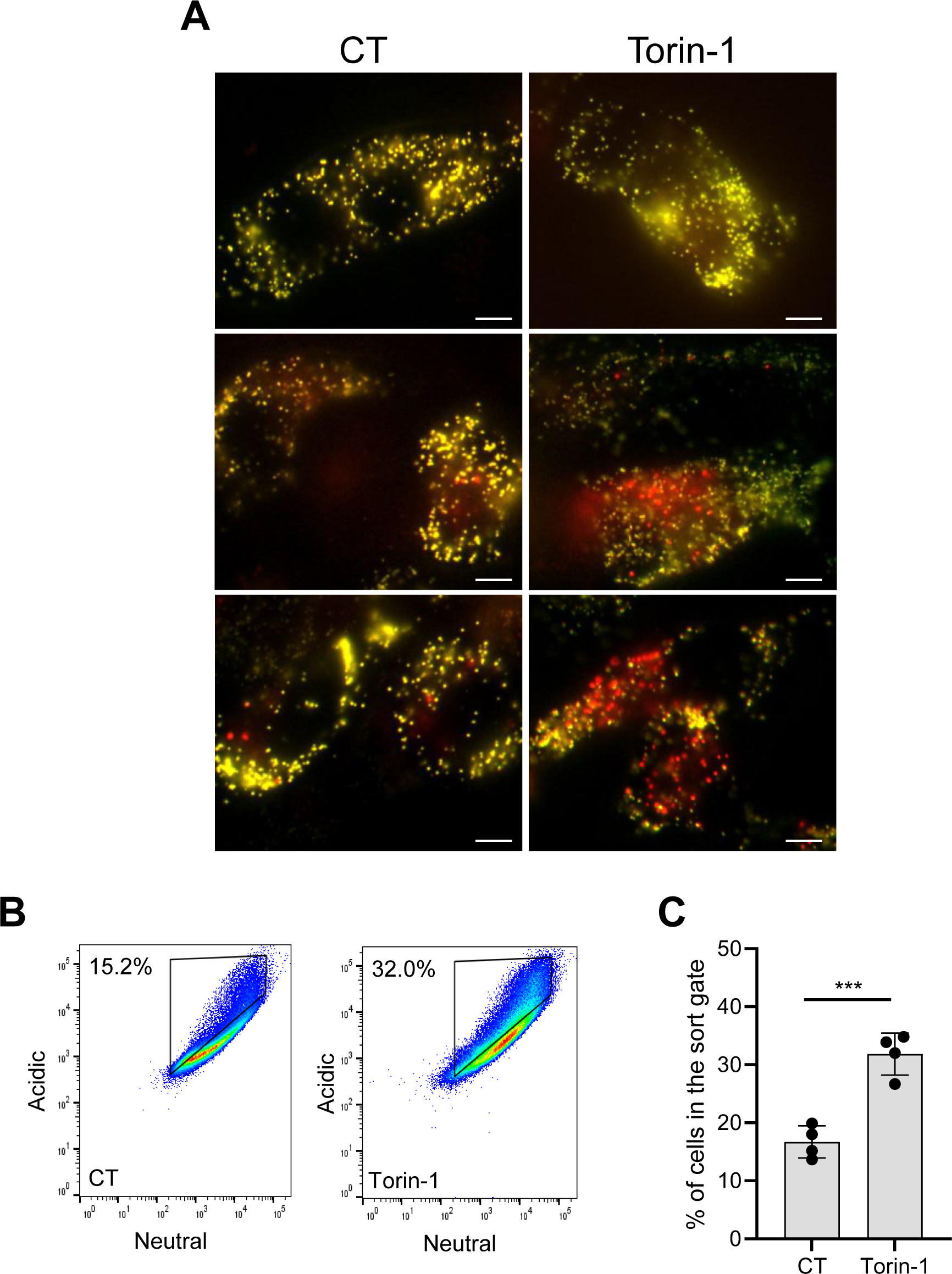
Torin-1 induces pexophagy in Hela cells lacking classical autophagy receptors. Po-mKeima pentaKO HeLa cells, deficient in NDP52, OPTN, TAX1BP1, NBR1, and SQSTM1 were treated overnight with 250 nM Torin-1 or left untreated (CT) and analyzed using fluorescence microscopy or flow cytometry. **(A)** Representative fluorescence overlay images of po-mKeima with excitation at 440 nm (false color: green) and 586 nm (false color: red) nm. Scale bar, 10 µm. **(B)** Representative flow cytometry plots for each group, with different colors indicating cell density at specific positions. **(C)** Quantification of the percentage of cells within the gated area. Data are presented as mean ± SD from 4 independent biological replicates. Statistical comparisons were performed (***, p < 0.001).

## Discussion

In this study, we identified PEX14 as a central mediator of OPTN-driven pexophagy. Domain mapping analyses revealed that OPTN variants lacking key autophagic motifs, specifically the ATG9A- and LC3-binding motifs, were still capable to initiate pexophagy when ectopically expressed. Notably, this induction was dependent on the NOA domain, a region known for its affinity for linear ubiquitin chains (Li et al., 2018) and its role in facilitating homotrimer formation (Gao et al., 2014). In addition, the OPTN_Δ474-479_ variant, which contains a NOA domain unable to bind polyubiquitin chains (Turturro et al., 2014), was still capable of triggering pexophagy. Conversely, knockdown of endogenous OPTN inhibited pexophagy induction by the OPTN_217-577_-GFP variant. These findings suggest that pexophagy induction by OPTN_217-577_-GFP likely occurs through oligomerization with endogenous OPTN. GFP-Trap pull-down experiments further revealed that the NOA domain of OPTN, rather than its ubiquitin-binding activity, is essential for its interaction with the coiled-coil domain of PEX14. Notably, previous research has shown that (i) the NOA domain can bind protein aggregates in a ubiquitin-independent manner (Korac et al., 2013), and (ii) the coiled-coil domain of PEX14 is a key determinant for homo-oligomerization (Fransen et al., 2002; Itoh and Fujiki, 2006).

Our findings collectively indicate that OPTN-mediated pexophagy is promoted through interactions with PEX14, a relationship further supported by the co-overexpression of OPTN and PEX14 in HeLa cells. Interestingly, inactivating classical autophagy receptors in these cells did not affect peroxisome levels or interfere with Torin-1-induced pexophagy. This suggests that neither OPTN nor other known pexophagy receptors, such as SQSTM1 and NBR1 (Deosaran et al., 20132; Yamashita et al., 2014; Zhang et al., 2015; Riccio et al., 2019), are essential for basal pexophagy under the tested conditions. This observation also supports the existence of non-canonical, ubiquitin-independent pexophagy pathways (Hara-Kuge and Fujiki, 2008; Jiang et al., 2015; Li et al., 2017; Wilhelm et al., 2022; Barone et al., 2023).

Consequently, this raises the possibility that canonical autophagy receptors may regulate pexophagy under specific conditions that are yet to be clarified.

Considering the established roles of phosphorylation, ubiquitylation, and acetylation in initiating, regulating, and fine-tuning autophagy (McEwan and Dikic, 2011), a plausible hypothesis for explaining cell type-dependent variations in OPTN-mediated pexophagy may involve cell type-specific posttranslational modifications. Notably, both OPTN and PEX14 are substrates for phosphorylation at multiple sites (see PTM sections in UniProt Q96CV9 and O75381, respectively). Specifically, OPTN phosphorylation by TBK1 (e.g., at S177 and S473) (Richter et al., 2016), a top-ranked proximity partner of OPTN in our BioID experiments, has been linked to selective autophagy processes, including the clearance of cytosolic bacteria (Wild et al., 2011), mutant protein aggregates (Korac et al., 2013), and damaged mitochondria (Richter et al., 2016). Although both OPTN_217-577_-GFP and OPTN_Δ474-479_-GFP can induce pexophagy, it remains unclear whether TBK1-mediated phosphorylation at S177 and S473 is essential for this process, particularly considering that these variants may oligomerize with endogenous OPTN. In addition, assuming that protein import into the peroxisomal membrane is a stochastic process, another intriguing hypothesis is that PEX14 concentrations in the membrane could act as a “timer” for peroxisome aging, with higher levels marking older peroxisomes that are more likely to be targeted for pexophagy. While testing these hypotheses was beyond the scope of the present study, they open interesting avenues for future research.

An important question is the role of OPTN-mediated pexophagy in normal physiology and disease. While much of the research has focused on autophagy receptors such as SQSTM1 and NBR1, the role of OPTN in pexophagy remains largely unexplored. Proteomics analysis of retinas from OPTN-E50K mice revealed that the KEGG peroxisome pathway was among the most significantly upregulated pathways in 18-month-old mice compared to control mice (Liu et al., 2021). This upregulation may result from the OPTN E50K mutation impairing the effective degradation of peroxisomes. In addition, increased expression of OPTN has been associated with elevated pro-inflammatory cytokine levels (Vittitow et al., 2002; Sudhakar et al., 2013), and their combined induction with pexophagy during prolonged fasting in catalase-deficient mouse livers (Dutta et al., 2021) suggests that this physiological upregulation of OPTN may drive pexophagy to meet metabolic and energetic demands. Furthermore, a recent study identified elevated pexophagy in Zellweger spectrum disorder patients with C-terminally truncated PEX14 variants (p.A196Lfs∗34 and p.A196Vfs∗34), a phenomenon only partially alleviated by NBR1 knockdown (Waterham et al.,2023). This finding suggests that truncation of PEX14 near its coiled-coil domain may expose this domain, thereby activating OPTN-mediated pexophagy in the patient cells. These findings position OPTN as a potential key regulator of pexophagy in both normal and pathological conditions.

A limitation of this study is the reliance on OPTN overexpression to reliably detect and quantify pexophagy. However, given that peroxisomes in cultured cells have a half-life of approximately two days (Huybrechts et al., 2009), while autophagosomes are rapidly turned over with an average lifespan of just 8 minutes (Papadopoulos and Pfeiffer, 1986), it is unsurprising that OPTN is infrequently observed on peroxisomes under basal conditions. This presents a challenge for studying OPTN-mediated pexophagy in native settings. This is similar to the difficulties encountered in studying the well-established PINK-Parkin-regulated mitophagy pathway, where many studies rely on cellular models involving parkin overexpression and chemicals or conditions that induce mitochondrial depolarization (Dhar et al., 2024). These considerations underscore the challenges of studying pexophagy and other selective autophagy pathways, such as PINK-Parkin-mediated mitophagy (McWilliams et al., 2018), in physiological conditions.

In summary, this study represents a foundational step in understanding OPTN-mediated pexophagy. Further research into the molecular mechanisms underlying OPTN-mediated pexophagy may enhance our knowledge of disorders associated with peroxisome dysfunction and OPTN dysregulation. Such insights may eventually contribute to the development of innovative therapeutic strategies for these conditions, presenting new opportunities for intervention in diseases linked to impaired selective autophagy.

## Materials and Methods

### DNA manipulations and plasmids

The mammalian expression plasmids encoding mCherry-PTS1 (pMF1218) (Nordgren et al., 2012), po-mKeima (pMF2005) (Li et al., 2023), non-tagged HsPEX14 (Will et al., 1999), HsPEX14-GFP (pMF120) (Fransen et al., 2004), GFP-LC3 (Nordgren et al., 2012), or GFP_Nb_-PM (pMF1968) (Walton et al., 2017) have been described elsewhere. The flippase (Flp) recombinase expression vector (pOG44; Thermo Fisher Scientific, V600520), the Flp-In^TM^ inducible expression vector (pcDNA5/FRT/TO; Thermo Fisher Scientific, V652020), the lentiviral GFP expression vector (pLenti CMV GFP Puro; Addgene, plasmid 17448), and the plasmids encoding GFP (Clontech, 6085-1), the pOPINE GFP nanobody (Addgene, plasmid 49172 (Kubala et al., 2010)), non-tagged OPTN_1-577_ (Origene, SC319257), OPTN_1-577_-GFP (Addgene, plasmid 27052 (Park et al., 2006)]), OPTN_1-209_-GFP (Addgene, plasmid 68839 (Turturro et al., 2014)), OPTN_217-577_-GFP (Addgene, plasmid 68841 (Turturro et al., 2014), or OPTN_D474N_-GFP (Addgene, plasmid 68847 (Turturro et al., 2014)) were commercially obtained. The miniTurbo BirA template was kindly provided by Dr. P. Kim (University of Toronto, Canada).

The following plasmids, annotated as “name (encoded protein; cloning vector; cloning procedure; template; forward primer (5’→3’); reverse primer (5’→3’); restriction enzymes)”, were specifically generated for this study: pDI1 (OPTN_Δ545-577_-GFP; Addgene 27052; Q5 Site-Directed Mutagenesis; Addgene 27052; gactggctgcatcattcgggatccaatgg; tgatgcagccagtccctgtcctcagc; not applicable (NA)), pDI2 (OPTN_Δ474-479_-GFP; Addgene 27052; Q5 Site-Directed Mutagenesis; Addgene 27052;

actgttctgcagcgagagagaaaattcatgagg; tcgctgcagaacagtaaacttccatctgagcc; NA), pHL30 (miniTurbo-OPTN_1-577_-GFP; Addgene 27052; cloning of digested PCR product; miniTurbo-BirA; cgagcgctagcaccatggcgatcccgc; cgagggaattcgcgacccacccgacccgcccttttcggcagaccgcag; NheI/EcoRI), pHL31 (OPTN_1-577_-GFP; pcDNA5/FRT/TO vector; cloning of digested PCR product; Addgene 27052; cgagggtaccaccatggctatgtcccatcaac; cgaggcggccgcttacttgtacagctcgtccatg; KpnI/NotI), pHL32 (OPTN_1-577_-GFP; pLenti CMV GFP Puro; cloning of digested of PCR product; Addgene 27052; gacaccgactctagaggatccaccatggctatgtcccatcaa; atggtggcgaccggtggatcccgaatgatgcaatccatcacg; BamHI), pHL34 (po-mKeima; pLenti CMV GFP Puro; cloning of digested PCR product; pMF2005; cgaggggatccaccatggtgagcgtgatcg; cgagggtcgacttacagcttgctcttgcccag; BamHI/SalI), pHL36 (OPTN_295-360_-GFP; pEGFP-N1; cloning of digested PCR product; Addgene 27052; gagaaaggcccggagactttagagctacaagtggaaagcatg; ccacctgtagctctaaagtctccgggcctttct; BglII/PstI), pHL37 (OPTN_422-455_-GFP; pEGFP-N1; cloning of digested PCR product; Addgene 27052; gaaaaagtggacagggcagaggacctggaaaccatg; catggtttccaggtcctctgccctgtccactttttc; BglII/PstI), pHL38 (OPTN_Δ454-520_-GFP; pEGFP-N1; cloning of digested PCR product; Addgene 27052; aagcaaaccattgccaagcatggggcgagaacaagtgac; cctggcaatggtttgcttcatttcatcc; BglII/PstI), pMF235 (PEX14_65-377_-GFP; pEGFP-N1; cloning of digested PCR product; pMF120;

gggaagatctatggccttccagcagtcggg; aatctgcaggtcccgctcactctcgtt; BglII/PstI); pMF415 (PEX14_1-313_-GFP; pEGFP-N1; cloning of digested PCR product; pMF120; gggaagatctatggcgtcctcggagcag; ggactgcagcacctccatccgcacctg; BglII/PstI), pMF663 (PEX14_Δ139-199_-GFP; pEGFP-N1; cloning of digested fusion PCR product; pMF120; primer set 1: gggaagatctatggcgtcctcggagcag and gatccagttggtgccgcccaggatgag; primer set 2: atcctgggcggcaccaactggatcctggag and aatctgcaggtcccgctcactctcgt; BglII/PstI), and pMF682 (PEX14_Δ110-126_-GFP; pEGFP-N1; cloning of digested fusion PCR product; pMF120; (primer set 1: gggaagatctatggcgtcctcggagcag and gtatttcttgtagtaatctcgccatcggga; primer set 2: tggcgagattactacaagaaatacctgctc and aatctgcaggtcccgctcactctcgtt; BglII/PstI). For the construction of pHL34, pHL36, pHL37, and pHL38, the PCR products were treated with DpnI and then inserted into the cloning vector using the NEBuilder HiFi DNA Assembly cloning kit. The primers were obtained from Integrated DNA Technologies. The *TOP10F’ E. coli* strain (Thermo Fisher Scientific, C3030-06) was routinely used as a cloning and plasmid amplification host. However, upon using the NEBuilder® HiFi DNA Assembly Master Mix (New England Biolabs, M5520AVIAL) or the Q5 Site-Directed Mutagenesis Kit (New England Biolabs, E0554), the DpnI (Thermo Fisher Scientific, ER1707)-treated mixtures, free of parental plasmid, were transformed in DH5α cells (Thermo Fisher Scientific, 18265017).

All plasmids were validated by DNA sequencing (LGC Genomics).

### Cell lines and culture conditions

The following cell lines were used for this study: human embryonic kidney cells (HEK-293, ECACC 85120602), Flp-in T-REx 293 cells (Thermo Fisher Scientific, R78007), lenti-GFP Flp-In T-REx 293 cells (this study), lenti-OPTN_1-577_-GFP Flp-In T-REx 293 cells (this study), GFP Flp-In T-REx 293 cells (this study), OPTN_1-577_-GFP Flp-In T-REx 293 cells (this study), lenti-po-mKeima Flp-In-T-REx 293 cells (Li et al., 2023), human cervical cancer cells (HeLa, ECACC 93021013), po-mKeima HeLa cells (Li et al., 2023), HeLa cells lacking NDP52, OPTN, TAX1BP1, NBR1, and p62 (pentaKO) (Lazarou et al., 2015), po-mKeima pentaKO (this study), human colorectal carcinoma cells (HCT116, ECACC 91091005), and po-mKeima HCT116 cells (this study). The novel Flp-In T-REx 293, lenti-Flp-In T-REx 293, and po-mKeima-expressing cell lines were generated as detailed elsewhere (Li et al., 2023; Lismont et al., 2019; Campeau et al., 2009; Bhalla et al., 2024). HEK-293, Flp-in T-REx 293, and HeLa cells were cultured in DMEM medium (Gibco, 31966-047) supplemented with 10% (v/v) fetal bovine serum (FBS; Biowest, S181B) and 0.2% (v/v) MycoZap (Lonza, VZA-2012) in a humidified atmosphere of 5% CO_2_ at 37°C. HCT116 cells were cultured similarly, using DMEM/F12 medium (Gibco, 21041-033) and the same supplements plus 2 mM UltraGlutamine I (Lonza, BE17-605E/U1).

### Cell transfection and treatment procedures

Flp-In T-REx 293 cells were either electroporated with the Neon Transfection system (Thermo Fisher Scientific) (Li et al., 2023) for fluorescence microscopy imaging and flow cytometry analyses or transfected using the JetPrime reagent (Polyplus, 114-01) for IP experiments. HCT116 and HeLa cells were transfected with the Lipofectamine 3000 reagent (Invitrogen, L3000015). To prevent autophagosome-lysosome fusion, the cells were pretreated for 2 h with 200 nM bafilomycin A1 (BafA1; Cayman Chemical, 11038). The same volume of DMSO (MP Biomedicals, 02196055) was included as vehicle control. To induce autophagy, the cells were treated overnight with 250 nM Torin-1 (Tocris, 4247) or an equivalent volume of vehicle control (DMSO). To image mitochondria, the endoplasmic reticulum, and lysosomes, the cells were incubated with MitoTracker™ Red CM-H2Xros (Thermo Fisher Scientific, M7513; 1:2000), ER-Tracker™ Blue-White DPX (Thermo Fisher Scientific, E12353; 1:2000), and LysoTracker™ Red DND-99 (Thermo Fisher Scientific, L7528; 1:5000), respectively. To downregulate OPTN expression, the cells underwent transfection (using Lipofectamine 3000) twice with the OPTN DsiRNA hs.Ri.OPTN.13.10 (sense strand: 5’-aagcuaaauaaucaagc-3’; antisense strand: 5’-cuuucauggcuugauuau-3’; final concentration: 50 nM), with a 48-h interval between each transfection. A non-targeting DsiRNA (Integrated DNA Technologies, 51-01-14-03), which does not recognize any human sequences, was used as a control.

### Antibodies and streptavidin conjugates

For immunofluorescence (IF) or immunoblotting (IB), the following primary antibodies were used: rabbit anti-LC3B (CST, 2775S; IB, 1:1000); rabbit anti-OPTN (Abcam, ab23666; IF, 1:50; IB, 1:1000), rabbit anti-SQSTM1 (Abcam, ab109012; IB, 1:20000), mouse anti-GAPDH (Sigma-Aldrich, G8795; IB, 1:2000), rabbit anti-GFP (IB, 1:50000) (Fransen et al., 2001), rabbit anti-PEX13 (IB: 1:1000) (Fransen et al., 2001), rabbit anti-PEX14 (IB, 1:1000) (Amery et al., 2000), and mouse anti-PEX14 (IF, 1:1000) (Van Veldhoven et al., 2020). The secondary antibodies for IF were conjugated to Alexa Fluor 488 (Invitrogen, A11017; 1:2000), Texas Red (Calbiochem, 401355; 1:200), or CF^TM^ 350 (Sigma-Aldrich, SAB4600406; 1:500). Streptavidin-horseradish peroxidase (IB, 1:3000) was from Sigma-Aldrich (RABHRP3).

### Immunoblotting and densitometric analysis

Antigen-antibody and biotin-streptavidin complexes were detected by using the Amersham ECL Western Blotting Detection Reagent (Cytiva, RPN2235) in combination with the ImageQuant LAS4000 system (GE Healthcare). Images were further processed and quantified with AlphaView Software (ProteinSample) or ImageJ software (Schneider et al., 2012).

### Fluorescence microscopy

Fluorescence microscopy was conducted essentially as previously described (Ramazani et al., 2021). Briefly, fluorescence was assessed using a motorized inverted IX-81 microscope (Olympus) controlled by cellSens Dimension software (version 2). The microscope was equipped with (i) a temperature, humidity, and CO_2_-controlled incubation chamber, (ii) a CoolLED pE-4000 illumination system, (iii) a 100x Super Apochromat oil immersion objective, (iv) the filter cubes F440 (excitation: 422–432 nm; dichroic mirror: 600 nm; emission: 610 long pass), F482 (excitation: 470–495 nm; dichroic mirror: 505 nm; emission: 510–550 nm), and F562 (excitation: 545–580 nm; dichroic mirror: 600 nm; emission: 610 nm long pass). For live-cell imaging, cells were seeded and imaged in FluoroDish cell culture dishes (World Precision Instruments, FD35), either pre-coated with 25 μg/mL polyethyleneimine (MP Biomedicals, 195444) for HEK-293 cells or left uncoated for other cell types. For immunofluorescence microscopy, cells were seeded onto (polyethyleneimine-coated) glass coverslips, and subsequently fixed and processed following methods outlined elsewhere (Huybrechts et al., 2009). Nuclei were counterstained with 6-diamidino-2-phenylindole (DAPI; Roche, 10236276001). Image acquisition and analysis were performed using cellSens Dimension software (version 2.1) (Olympus). Nanobody-based plasma membrane translocation assays were carried out according to established procedures (Walton et al., 2017).

### Structured illumination microscopy

Cells were cultured and imaged in Fluorodishes (World Precision Instruments, FD35). For image acquisition, a Nikon Ti2 N-structured illumination (SIM) S microscope (Nikon Instruments) equipped with an ORCA-Flash4.0 sCMOS camera (Hamamatsu, C13440) and a Nikon SR Apo TIRF 100x objective lens (NA 1.49) was controlled by Nikon NIS-Elements imaging software (version 5.21.03). Multicolored images were acquired in three-dimensional SIM mode (with three angles and five phases) with a Z-stack (0.100 µm step size) at room temperature. The fluorescence of GFP, excited at 488 nm, was captured using a 525/30 bandpass filter, and the fluorescence of mCherry, excited at 561 nm, was detected with a 595/30 bandpass filter. The images were reconstructed and analyzed using Arivis Vision4D (version 3.3) and Nikon NIS-Elements AR 5.40 software.

### Flow cytometry analysis

The po-mKeima-expressing cells were washed with DPBS and dislodged from the plate by trypsinization. Next, the cells were pelleted (300× g, 1 min), resuspended in culture medium, and analyzed through FACSymphony™ A1 flow cytometry (BD Biosciences) by using 405- and 561-nm lasers in combination with a 600–620 nm emission filter (for mKeima), and a 488-nm laser with a 505-560 nm emission filter (for GFP); For each sample, 100,000 events were collected and analyzed using BD FACSDiva v8.0.1 software (BD Biosciences).

### MiniTurbo proximity labeling, affinity purification, and sample preparation for LC-MS analysis

Cells were seeded in 15-cm dishes and transfected with 6 µg of a plasmid encoding miniTurbo-OPTN-GFP (pHL30). After 48 h, the cells were pretreated with 200 nM of BafA1 for 2 h and subsequently incubated with or without 50 µM D-biotin (Sigma-Aldrich, B4501) for 30 min. Next, the cells were rinsed with ice-cold PBS (Lonza, BE17-512F) and lysed on ice for 20 min using lysis buffer (50 mM Tris-HCl (pH 7.5), 150 mM NaCl, 1% (w/v) Triton X-100, 1 mM EDTA) supplemented with protease (Sigma-Aldrich, P2714; AppliChem, A0999) and phosphatase (Sigma-Aldrich, P5726) inhibitor cocktails. Subsequently, the lysates were sonicated (Diagnode Bioruptor UCD-200) in three 10-s cycles at high intensity and then cleared by centrifugation at 10,000 × g for 10 min. A small portion of the supernatant was taken for biotin IB validation, and the remaining supernatant was further incubated with 25 μL of Streptavidin Sepharose beads (Cytiva, 90100484) overnight at 4°C. After discarding the supernatant, the beads were washed 3 times with 200 μL of lysis buffer and 3 times with 200 μL of ammonium bicarbonate (50 mM, pH 7.8). Thereafter, the beads were resuspended in 200 μL of ammonium bicarbonate solution, and 250 ng of mass spectrometry grade trypsin (Promega, V528A) was added. The samples were digested for 14 hours at 37°C in an Eppendorf Thermomixer C (speed: 750 rpm), and a 2-μL sample was taken for tryptic digestion quality control using a Dionex™ UltiMate™ 3000 RSLC Nano System (Thermo Fisher Scientific). The reduction and carbamidomethylation of disulfide bridges in the supernatant were performed by treating the samples with 10 mM dithiothreitol (VWR, M109) for 30 minutes at 56°C, followed by the addition of 30 mM iodoacetamide (Sigma-Aldrich, I1149) for an additional 30 minutes at room temperature. Next, the carbamidomethylated samples were acidified using 10% trifluoroacetic acid (Biosolve, 202341) and subsequently subjected to desalting using Poros Oligo R3 reversed-phase resin (Thermo Fisher Scientific, 1133903). The bound peptides were washed twice with 0.1% formic acid (FA; Thermo Fisher Scientific, 28905) in water and eluted with 100 μL of 60% (v/v) acetonitrile (ACN; Sigma, 34851) in water containing 0.1% FA. Finally, the eluates were dried under vacuum, and the peptides were resolubilized in 10 μL of 0.1% FA.

### LC-MS analysis

Digested samples (10% of total volume) were injected into an UltiMate 3000 RSLC nano System (Dionex) coupled to an Orbitrap Lumos Tribrid Mass Spectrometer (Thermo Fisher Scientific). After initial loading, peptides were concentrated on a 75 μm × 2 cm C18 pre-column using 0.1% (v/v) trifluoroacetic acid at a flow rate of 20 µL/min. Sample separation was accomplished on a reversed-phase column (Acclaim C18 PepMap100; pore size: 75 µm; length: 50 cm) at 60°C using a binary gradient (A: 0.1% FA; B: 84% ACN with 0.1% FA) at a flow rate of 250 nL/min: 3% solvent B for 5 min, a linear increase of solvent B to 32% for 95 min followed by washing with 95% solvent B for 3 min and a linear decrease of solvent B to 3% for 1 min. Peptides were ionized by using a nanospray ESI source. MS survey scans were acquired on the Orbitrap Lumos Tribrid by using the following settings: mass spectrometer was operated in data-dependent acquisition mode (DDA) with full MS scans from 350 to 1500 m/z at a resolution of 120,000 at 200 m/z (Orbitrap) using the polysiloxane ion at 445.12002 m/z as lock mass. The automatic gain control (AGC) was set to 4E5, the maximum injection time to 100 ms, and the RF amplitude to 40%. The top intense ions (ion intensity ≥ 5E3, charge state = 2-7) were selected within a 1.2 m/z mass isolation window for fragmentation at a normalized collision energy of 32% in each cycle 3 seconds of the acquisition analysis, following each survey scan. The dynamic exclusion time was set to 60 s. Fragment ions were acquired in the ion trap.

### Data analysis of miniTurbo proximity labeling

Raw data were searched against the UniProt human database (June 2023; 20,409 target sequences, including OPTN-GFP) using Sequest HT on Proteome Discoverer v2.5. Precursor mass tolerance was limited to 20 ppm and fragment mass tolerance to 0.5 Da. Cleavage specificity was set to fully tryptic, allowing for a maximum of two missed cleavages.

Carbamidomethylation of cysteines (+57.0214 Da) was defined as a fixed modification and oxidation of methionine (+15.9949 Da) as a variable modification for all searches. The results were evaluated with Percolator for false discovery rate (FDR) estimation and data were filtered at ≤1% FDR on the PSM and peptide level and filtered ‘master’ proteins at protein level (Käll et al., 2007). Unique peptides (except modified peptides) were taken for Label free quantification. Next, the sum of top three peptide intensities was calculated to determine the protein intensities. Protein intensities were normalized to total peptide amount. P-values between the sample groups (four biological replicates, with or without D-biotin treatment) were determined by a two-sided t-test with FDR correction. Common contaminating proteins were removed from the hit lists. The significance cut-offs employed were unique peptides ≥ 2, adjust p-value ≤ 0.05 and log_2_(ratio biotin/no biotin) ≥ 1. Label free quantification data were plotted and visualized by GraphPad Prism 9.0.0.

### Immunoprecipitation using GFP-Trap

HEK-293 cells were cultured in 10-cm cell culture dishes and transfected with 6 μg of plasmid. Two days later, the cells were cultured for 2 h in the presence of 200 nM BafA1. After rinsing with ice-cold PBS, cleared supernatant fractions were prepared similarly to the miniTurbo proximity labeling samples and subjected to a GFP-Trap assay (Claes et al., 2023). Briefly, the supernatants were divided into two portions for input detection and GFP-Trap.

The input fractions were supplemented with 3x SDS-PAGE sample buffer for IB analysis and boiled at 100 °C for 10 min. The supernatants for GFP-Trap immunoprecipitation were pre-cleared with 30 μL of homemade BSA beads for 30 min at 4 °C, followed by incubation with 25 µL of homemade GFP-Trap beads for 2 h at 4 °C (the BSA- and pOpine-GFP nanobody-conjugated beads were prepared as described elsewhere (Rothbauer et al., 2008)). Next, the beads were washed 6-8 times with high saline lysis buffer (150 mM Tris-HCl (pH 7.5), 500 mM NaCl, 1% (w/v) Triton X-100, 1 mM EDTA), boiled in 3x SDS-PAGE sample buffer, and after a quick spin, the supernatants were subjected to IB (Li et al., 2023).

### Statistical analysis

Statistical analysis involved conducting one-way ANOVA followed by Tukey’s post hoc test for multiple comparisons. Student’s unpaired two-tailed t-test was used to compare two groups. All statistical analyses were conducted using GraphPad Prism (version 9.0.0 for Windows 64-bit, GraphPad Software, San Diego, CA, USA). Results with a p-value less than 0.05 were considered statistically significant.

### Varia

The data used to generate Fig. 3A, the schematic depicting the distinct domains and major interactions of OPTN, were gathered from multiple sources (Weil et al., 2018; Zhao et al., 2023; Kachaner et al., 2012; Swarup and Sayyad, 2018). The manuscript’s language was refined with the assistance of ChatGPT.

### Supplemental material

The following supporting information is available: Figure S1: Expression of OPTN-GFP triggers pexophagy in HCT116 but not in HeLa cells; Figure S2: Impact of OPTN-GFP expression on organelle morphology in HEK-293 cells; Figure S3: OPTN-mediated pexophagy can be inhibited by bafilomycin A1 and does not affect general autophagy; Figure S4: Analysis of the OPTN-PEX14 interaction using GFP-trapping; Figure S5: Validation of the effectiveness of the OPTN DsiRNA.

### Data availability statement

The raw data from the LC-MS runs can be accessed via ProteomeXchange (PRIDE database) with the identifier PXD053489 (reviewers can use the username and the password “bqCvvzGODy8Z”). The data used to generate Figure 3B are available on FigShare (https://figshare.com/s/1e6bbf0320d72825eebb). All other data supporting the findings of this study are provided within the article or its supplementary materials.

## Acknowledgments

The authors acknowledge the following entities and individuals for their invaluable support: the VIB Bio Imaging and FACS Core facilities (KU Leuven) for assisting in super-resolution imaging and FACS analysis, the John Creemers lab (KU Leuven) for lentiviral packaging, Dr. R. Youle (National Institutes of Health, Bethesda, MD, USA) for the pentaKO HeLa cell line, Dr. M. Bollen (KU Leuven, Belgium) for the HCT116 cell line, Ms. S. Lemaire and Mr. X.Y Cao (KU Leuven, Belgium) for valuable advice regarding the GFP-Trap assay, Dr. R.J. Wanders (University of Amsterdam, The Netherlands) for the plasmid encoding non-tagged HsPEX14, Dr. P. Kim (University of Toronto, Canada) for the miniTurbo BirA template, and Dr. B. Yue (University of Illinois, Chicago, IL, USA) for the plasmids encoding OPTN_1-577_-GFP, OPTN_1-209_-GFP, OPTN_211-577_-GFP, or OPTN_D474N_-GFP. This work received support from the Research Foundation-Flanders (G091819N, G0A8619N, 1213623N), the China Scholarship Council (201906790005), KU Leuven (C14/18/088), and the Ministry of Higher Education of the Arab Republic of Egypt.

## Author contributions

H.L. and M.F. conceptualized the study and designed the experiments. H.L, S.C., and S.V. conducted the proteomics analyses. Y.L. generated the lentivirus particles and the stable lentiviral cell lines. D.I. and W.V generated the plasmids encoding OPTN_Δ474-479_ and OPTN_Δ545-577_. H.L., B.V., and S.C performed the other experiments. H.L, C.L., C.C., M.A.H., M.B., J.A., and M.F. analyzed and interpreted the data. C.L. and M.F. coordinated and supervised the project. M.F. and M.B. provided resources and funding for the investigation. H.L. and M.F. wrote the original draft of the manuscript, and all authors edited the draft and agreed on the content of the submitted version of the manuscript.

## Abbreviations

Baf A1: bafilomycin A1
BioID: biotin identification
CC: coiled-coil
CT: control
DsiRNA: Dicer-substrate short interfering RNA
FACS: fluorescence-activated cell sorting
FDR: false discovery rate
IB: immunoblotting
IF: immunofluorescence
LIR: LC3-interacting region
NOA: NEMO-OPTN-ABIN domain
PEX: peroxin
PTS1: C-terminal peroxisomal targeting signal 1
RR: relative ratios
ZF: zinc finger.

## Supplementary figure legends

**Figure S1.**
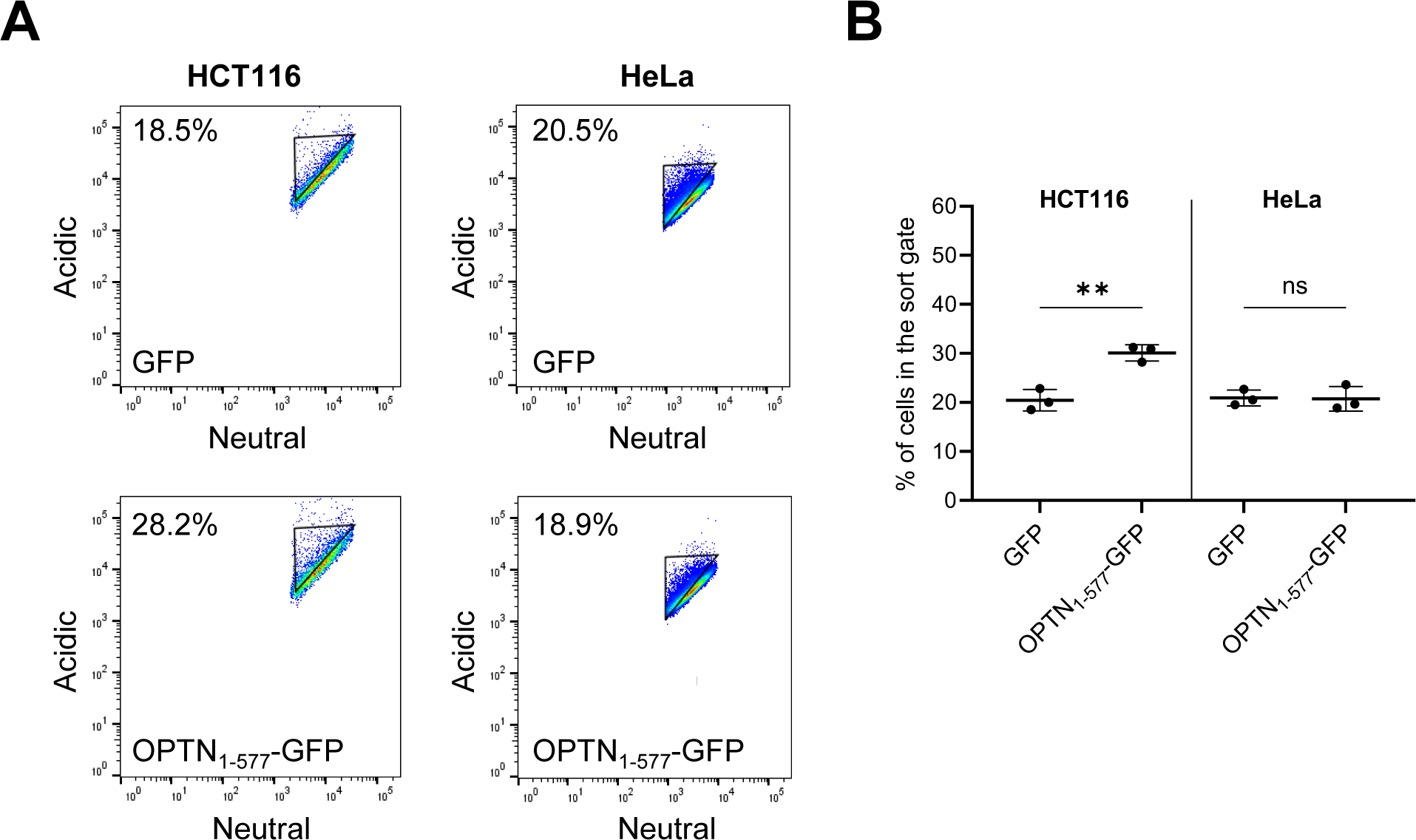
Expression of OPTN-GFP triggers pexophagy in HCT116 but not in HeLa cells. HCT116 and HeLa cells stably expressing po-mKeima were transfected with a plasmid encoding GFP or OPTN-GFP and cultured as described in the Materials and Methods section. After two days, the cells were harvested and processed for FACS analysis. To measure the percentage of cells undergoing pexophagy, single-cell GFP-positive populations were gated for a decrease in fluorescence intensity in the neutral channel. **(A)** Representative flow cytometry plots of each group (n = 3). The different colors represent the cell density at a given position. **(B)** Quantification of the percentage of cells in the gated area. The data are shown as the mean ± SD and represent the values of three independent biological replicates. All conditions were statistically compared to the GFP condition (ns, non-significant; **, p < 0.01).

**Figure S2.**
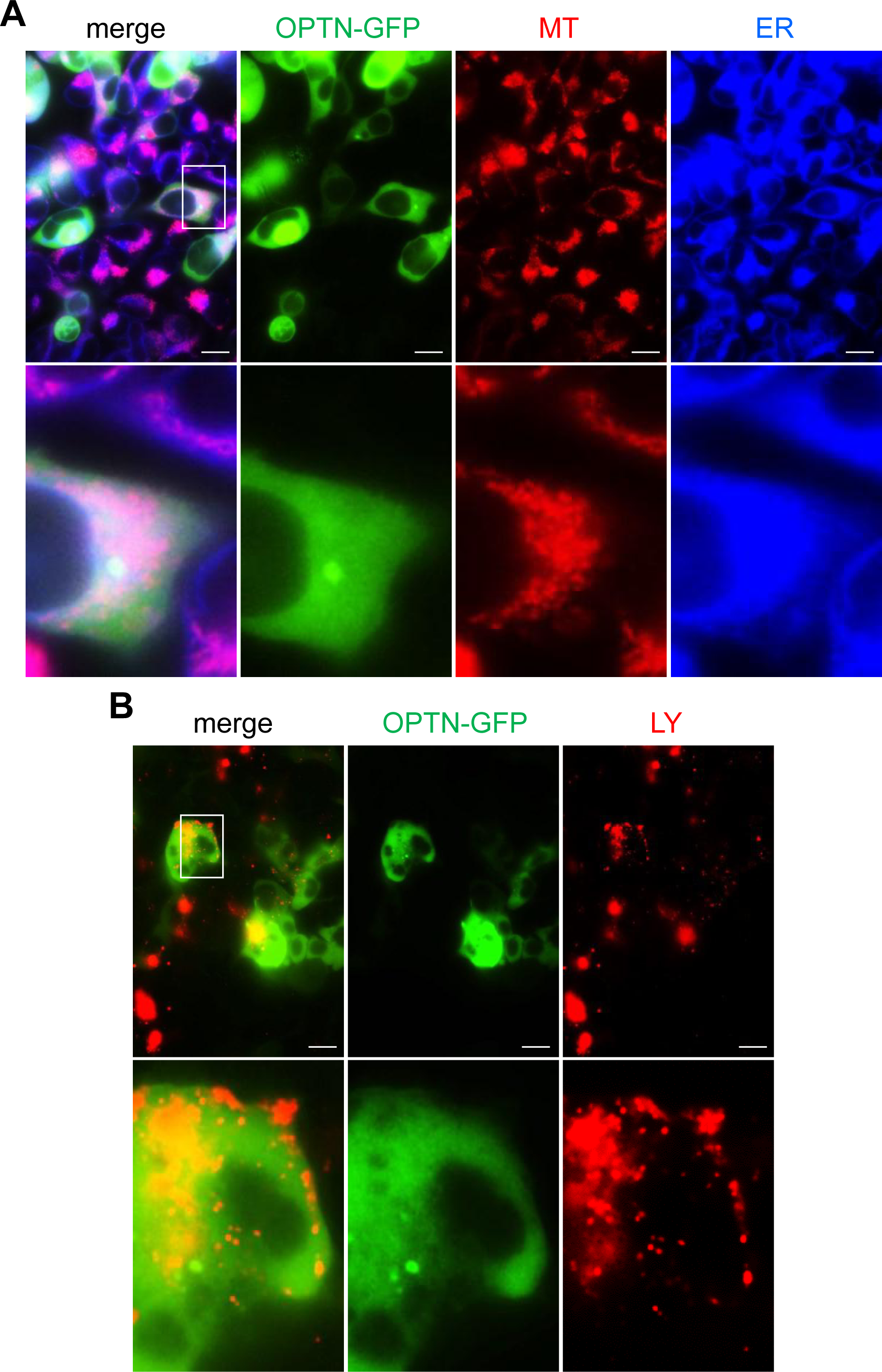
Impact of OPTN-GFP expression on organelle morphology in HEK-293 cells. **(A)** MitoTracker™ Red CM-H2Xros (MT) and ER-Tracker™ Blue-White DPX (ER) staining. LysoTracker™ Red DND-99 (LY) staining. Before live cell imaging, cells were incubated for 2 h with 200 nM BafA1. Representative images are shown. The lower panels depict enlargements of the boxed areas in the upper panels. Scale bars, 10 µm.

**Figure S3.**
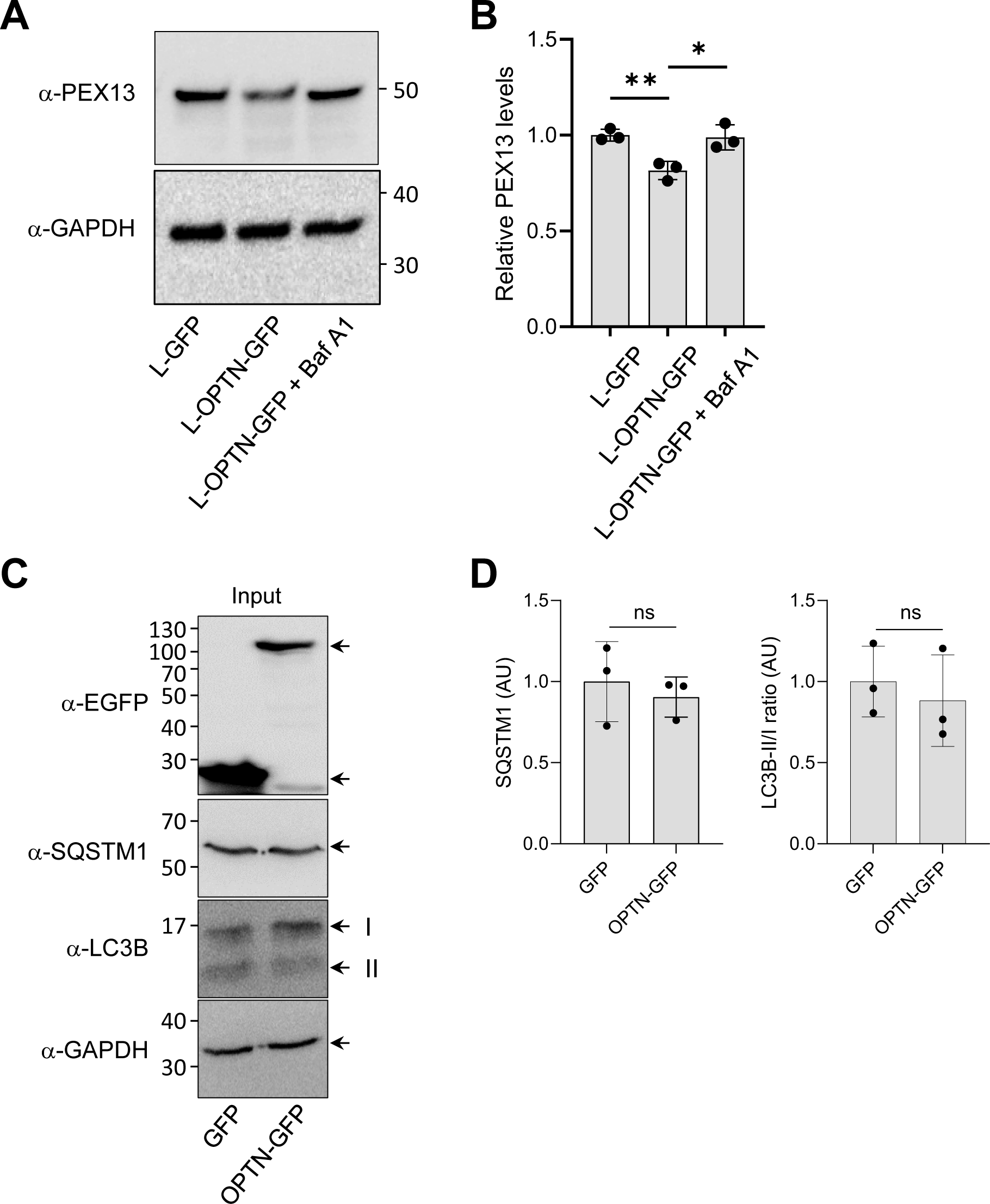
OPTN-mediated pexophagy can be inhibited by bafilomycin A1 and does not affect general autophagy. **(A,B)** Lenti-GFP Flp-In T-REx 293 (L-GFP) and Lenti-OPTN_1-577_-GFP Flp-In T-REx 293 (L-OPTN-GFP) cells were transfected with a plasmid encoding non-tagged PEX14. After 48 hours, the cells were incubated overnight in medium with or without 20 nM Bafilomycin A1 (Baf A1) and subsequently processed for immunoblotting using anti-PEX13 and anti-GAPDH antibodies. **(A)** Representative immunoblot for PEX13 and GAPDH. **(B)** Densitometry quantification of the PEX13 levels, with signal intensities normalized to the mean value of the L-GFP cells. Data represent the mean ± the standard deviation of three biological replicates (*, p < 0.05; **, p < 0.01). **(C,D)** HEK-293 cells were transiently transfected with plasmids encoding GFP or OPTN-GFP. After two days, cells were subjected to immunoblotting using antibodies against GFP, SQSTM1, LC3B, and GAPDH. (**C**) Representative blots are shown. **(D)** Densitometry quantifications are presented as means ± SD (n = 3), with data normalized to the average value of the GFP condition. Statistical analysis indicates no significant differences (ns).

**Figure S4.**
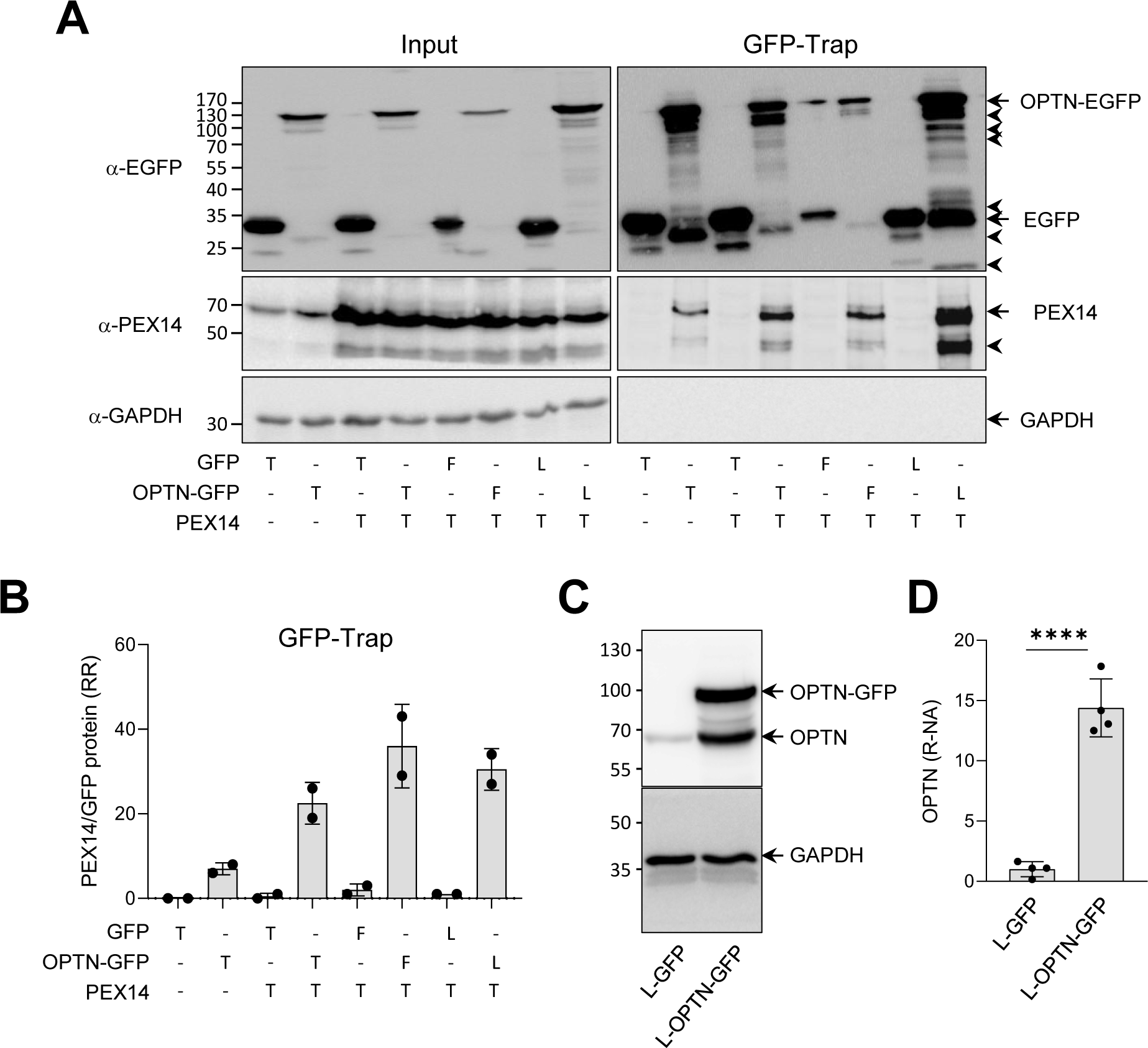
Analysis of the OPTN-PEX14 interaction using GFP-trapping. **(A)** Immunoblot analysis of input and GFP-Trap pull-down fractions from Flp-In T-REx 293 cells expressing the specified proteins. Representative images are shown. Specific protein bands and their degradation products are marked by arrows and arrowheads, respectively. Molecular mass markers (in kDa) are indicated on the left. **(B)** Densitometry quantifications of the relative ratios (RR) of PEX14/GFP and PEX14/OPTN-GFP retained on the GFP-Trap affinity matrix. The total signal intensities of the GFP proteins and PEX14 were both standardized to 100%. The bars represent the mean of two biological replicates. T, transient transfection; F, Flp-In integration; L, lentiviral transduction. **(C,D)** Relative expression of GFP-tagged OPTN compared to endogenous OPTN in lentiviral-transduced Flp-In T-REx 293 cells. **(C)** Total cell lysates from lentiviral (L-) transduced Flp-In T-REx 293 cells, which constitutively express either GFP or OPTN-GFP, were subjected to SDS-PAGE followed by immunoblotting using antibodies targeting OPTN and GAPDH.A representative image is shown, with specific protein bands indicated by arrows. Note that the 66 kDa band in the OPTN-GFP lane may represent either a degradation product of OPTN-GFP or endogenous OPTN stabilized by overexpression of OPTN-GFP. Molecular mass markers (in kDa) are indicated on the left. **(D)** Densitometry quantifications of the relative normalized amounts (R-NA) of OPTN. The expression levels were normalized to GAPDH, with the amount of OPTN detected in the GFP cells serving as the reference. The amount in the L-OPTN-GFP cells represents the sum of the 66 and 95 kDa bands. Bars represent the mean ± the standard deviation of four biological replicates (****, p < 0.0001).

**Figure S5.**
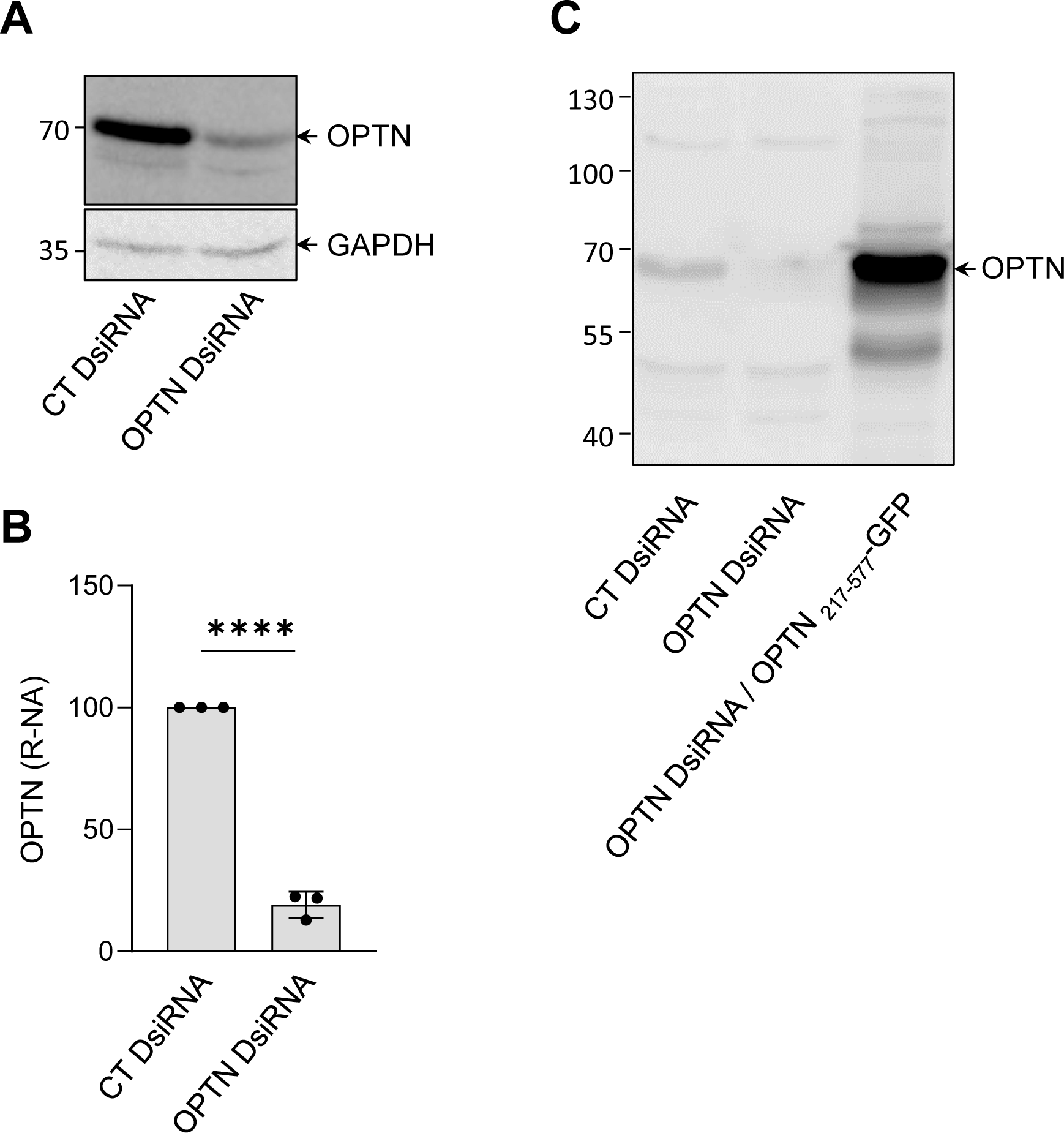
Validation of the effectiveness of the OPTN DsiRNA. Flp-In T-REx 293 cells stably expressing po-mKeima were transfected with either a control (CT) or OPTN DsiRNA and seeded in 6-well plates. On day two, the cells were transfected or not with a plasmid encoding OPTN_217-577_-GFP. On day three, the cells were transfected again with either DsiRNA CT or DsiRNA OPTN. On day four, the cells were harvested and processed for immunoblotting with antisera raised against the indicated proteins. **(A)** Representative blots showing the effectiveness of the OPTN DsiRNA. Specific protein bands are marked by arrows. **(B)** Densitometry quantifications of the relative normalized amounts (R-NA) of the OPTN levels shown in panel a (n = 3). Bars represent the mean ± the standard deviation of three biological replicates (****, p < 0.0001). **(C)** Comparison of the migration behavior of endogenous OPTN (CT DsiRNA condition) and OPTN_217-577_-GFP (OPTN DsiRNA / OPTN_217-577_-GFP condition). Molecular mass markers (in kDa) are indicated on the left.

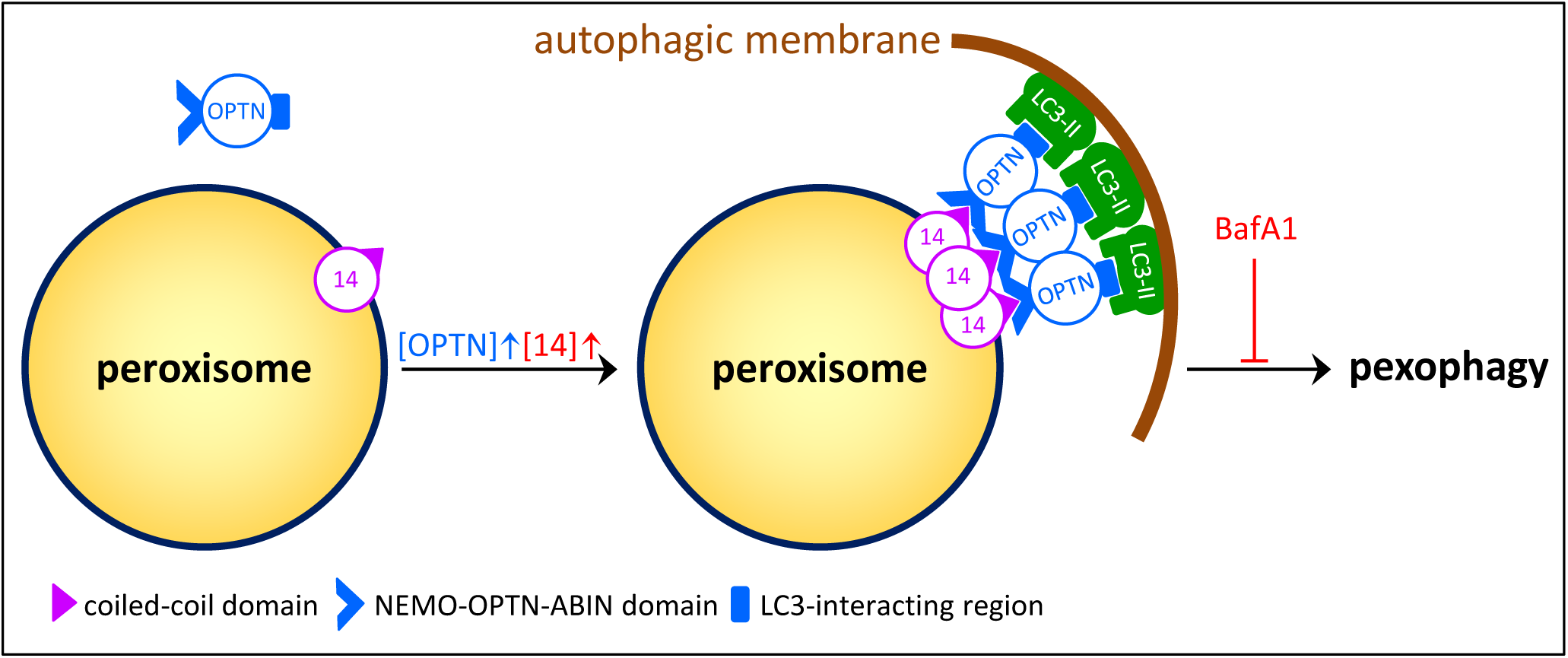

